# Evolutionary Constraints on RNA Polymerase Gene Positioning in the Genome of Fast-Growing Bacteria

**DOI:** 10.1101/2025.04.14.645983

**Authors:** Leticia Larotonda, Elisa Ojeda, María Belén Bordignon, Fabiana Fulgenzi, Marie-Eve Val, Diego Comerci, Briardo Llorente, Céline Loot, Didier Mazel, Alfonso Soler-Bistué

**Affiliations:** Laboratorio de Genómica Experimental Bacteriana, Instituto de Investigaciones Biotecnológicas (IIB), Escuela de Bio y Nanotecnologías (EByN), Universidad Nacional de San Martín (UNSAM) – Consejo Nacional de Investigaciones Científicas y Técnicas (CONICET), San Martín, Buenos Aires B1650, Argentina; Institut Pasteur, Université de Paris Cité, CNRS UMR 3525, Unité Plasticité du Génome Bactérien, Paris F-75015, France; Australian Research Council Centre of Excellence in Synthetic Biology, School of Natural Sciences, Macquarie University, Sydney, NSW, 2109, Australia; Australian Genome Foundry, Sydney, NSW, 2109, Australia

## Abstract

How gene order along chromosomes affects cellular homeostasis and genome evolution remains poorly understood. Bacterial chromosomes are organized along the replication origin (*oriC*)–terminus (*ter*) axis. The spatial arrengement of genes within this axis may influence cellular physiology, genome evolution, and transcriptional regulation. The catalytic core of the sole bacterial RNA polymerase (RNAP) is encoded by *rpoB* and *rpoC* within the universally conserved *rplKAJL-rpoBC* locus. In fast growing bacteria this locus is located near *oriC* potentially linking genome organization to cellular physiology. Here, we investigated the functional relevance of *rplKAJL-rpoBC* chromosomal positioning by relocating it within *Vibrio cholerae* genome. While viable, strains harboring *rplKAJL-rpoBC* far from *oriC* exhibited nutrient-dependent fitness defects. These phenotypes correlated with reduced locus copy number and RNAP abundance, without changes in its subcellular distribution. Introducing a second copy of *rpIKAJL-rpoBC* at a distal genomic location from *oriC* abolished the phenotypes, demonstrating that replication-associated gene dosage effects were the main mechanism behind the observed physiological changes. Further uncoupling *rpoB* and *rpoC* from the *rplKAJL-rpoBC* locus revealed that fitness costs were specifically linked to RNAP gene relocation. Our results reveal that selective pressures act to maintain RNAP-encoding genes near *oriC* to optimize replication-associated dosage effects, ensuring efficient RNAP production during exponential growth. We discuss the relationship between gene order and ecological strategies linked to bacterial growth. Our work underscores gene order as an underestimated but critical factor shaping cellular physiology and genome evolution.

**Significance:** Bacterial chromosomes have a highly plastic gene content but the importance of their spatial organization and gene order is started to be appreciated as a driver of cellular physiology. Growing evidence suggests that the spatial organization of bacterial genomes plays a critical role in gene expression, replication dynamics, and fitness. In this study, we demonstrate that the conserved positioning of *rpoB* and *rpoC*, encoding the catalytic core of RNA polymerase (RNAP), near the replication origin (*oriC*) is essential for optimal bacterial growth. By relocating the *rplKAJL-rpoBC* locus in *Vibrio cholerae*, we show that its chromosomal position directly impacts nutrient-dependent fitness, RNAP abundance, and replication-associated gene dosage effects. These findings suggest that selective pressures maintain genes encoding RNAP near *oriC* to ensure efficient transcriptional output during exponential growth. More broadly, our study highlights gene order as an underappreciated factor in bacterial genome evolution, beyond its known roles in operon structure and transcriptional regulation. Given the fundamental role of RNAP, our results have implications for understanding bacterial ecological strategies, adaptation, and synthetic genome design. By integrating positional genomics with physiological studies, we provide a theoretical and experimental framework for investigating how genome architecture shapes cellular function.

## Introduction

Over the decades it has been established that genes encode functions that usually are revealed by mutation and complementation. However, other aspects of inheritance such as epigenetics, non-coding DNA and DNA structure have received much less attention. During the last decade, the amount of information in genome databases increased steadily^1,2^. This is particularly true for prokaryotes whose genomes are relatively more straight-forward to study and easier to analyze. From this data, it is possible to infer that bacterial chromosomes have a single replication origin (*oriC*) where DNA replication starts, finishing at the chromosome terminus (*ter*). The *ori-ter* axis constitutes an organizational landmark that arranges the chromosome into two equally sized replichores^3^ which, in turn, conditions many genome features^4–6^. For example, the replication machinery movement along the DNA molecule leads to strand-dependent GC-skew and to the location of essential highly expressed genes on the leading strand to avoid detrimental head-on collisions with the transcription machinery^3,7^.

Recent work shows that the distribution of genes along the *ori-ter* axis is not random. Highly expressed and essential genes display an *oriC*-proximal location^7^. Such positional bias could be linked to cell physiology and ecological strategies. Pioneering work by Eduardo Rochás team has provided insights into the location of genes involved in the flux of genetic information: ribosomal RNA (rRNA), tRNAs, ribosomal proteins (RP) and RNA polymerase (RNAP) genes are found in close proximity to *oriC* especially in fast-growing bacteria^8^.

Gene location biases were also noticed in Nucleoid Associated Proteins, RNAP modulators and topoisomerases^9^. Importantly, their position along the *ori-ter* axis correlated to their expression along with the succession of growth phases. This means that genes nearby *oriC* are expressed during exponential phase while those close to *ter* are induced during stationary phase or in stress situations. A recent preprint studying more than 700 bacterial species, found that more than a half of gene families have specific chromosomal locations^10^.

The biased genomic location of certain genes is being increasingly supported by bioinformatic approaches, however, functional and mechanistic validation from experimental data is scarce. Recent works have shown that the genomic location of specific genes is necessary for the proper functioning of processes such as sporulation, natural competence for transformation, and cell growth^11–13^.

The positional genetics approach consists in the systematic relocation of key genes using recombineering techniques to address the impact of their chromosomal location on gene function and on cell homeostasis. In previous works, we applied this experimental procedure to the *s10-spc*-α locus (S10), that encodes half of the ribosomal proteins (RP) in *Vibrio cholerae,* the etiological agent of cholera^14–17^. *V. cholerae,* with a doubling time (DT) of 17 minutes, is a fast-growing microorganism that presents a bipartite genome. The replication of the 1Mbp secondary chromosome (Chr2) starts upon achieving of two-thirds of the duplication of the 3.1 Mbp main chromosome^18^ (Chr1).The transcription and translation genes are almost exclusively harbored by the main chromosome showing a clear location bias towards the *oriC* ^8,19^. In *V. cholerae,* genome-wide relocation of S10 led to a reduction in growth rate (GR) as a function of S10-*oriC* distance proving the concept that GR can be rationally tuned by relocating key loci. We also observed a fitness and infection capacity reduction associated with the relocation of the major ribosome protein loci far from *oriC*^14,15^. Importantly, adaptative laboratory evolution showed that growth defects associated to S10 relocation could not be suppressed suggesting that its location impacts *V. cholerae* evolutionary trajectory ^17^.

Overall, the chromosomal organization around the *ori-ter* axis reflects the importance of replication-dependent genome organization in bacteria^20^. However, only a few gene groups and specific loci have been experimentally tested for their dependence on chromosomal location for proper function. In this regard, while we provided extensive information on RPs, bioinformatic^10,21,22^ and experimental^23,24^ studies now suggest that RNAP genes may have an even greater impact on cell physiology.

Bacteria possess a single RNAP whose core is composed of the subunits α, β, and β’. Since the α subunit is present in excess, RNAP activity is limited by the abundance of β and β’ which are encoded by *rpoB* and *rpoC,* respectively^23,25^. These genes play a central role in genome expression. In *V. cholerae,* the *rpoBC-*encoding locus is located close to the *oriC* of the main chromosome (*ori1*). Here, we investigated the physiological importance of its chromosomal positioning using a positional genetic approach. We showed that the genomic location of the RNAP-encoding locus impacts cell physiology and fitness. Through genetic dissection, we attributed the observed phenotypes to the positioning of *rpoB* and *rpoC*.

## Results

### The structure of the *rplKAJL-rpoBC* locus is widely conserved and its location is biased towards *ori1* in *Vibrionaceae*

The *rpoB* and *rpoC* genes are physically linked to translation-related genes within the *tuf-secE-nusG-rplKAJL-rpoBC* locus (*rplKAJL-rpoBC* locus, Figure 1). This locus also encodes four tRNAs, the translation elongation factor EF-Tu, a subunit of the type II secretion system, the transcription elongation factor Tuf, and four RPs. The RNAP core genes are part of a larger operon that is mainly transcribed from the promoters of *rplJ* and *rplK* genes^23,26,27^. The *rplKAJL-rpoBC* operon (Figure 1) has been reported to be widely conserved among bacteria, archaea and eukaryotic organelles^28,29^. A closer analysis of *rplKAJL-rpoBC* locus architecture confirms synteny among some bacterial models. However, some divergences exist. In *Bradyrhizobium diazoefficiens* USDA110 and *Actinomyces israelii,* some genes separate *rplKA* from *rplJL* (Figure S1). Also, in cyanobacteria and in plastids, the *rpoC* gene is divided into *rpoC1* and *rpoC2*^30^. Next, we examined the genomic location of the *rplKAJL-rpoBC* locus among 31 available genomes within the *Vibronaceae* family (Table S2). The *rplKAJL-rpoBC* locus is consistently located on the main chromosome and, on average, is positioned near *ori1* within the initial 12.5% of the *ori-ter* axis. The furthest distance between *ori1* and the *rplKAJL-rpoBC* is observed in *Vibrio nigripulchritudo* SFn1 where the locus is at 36% of the *ori-ter* axis. *V. cholerae* isolates display the locus either at 13% or at 22%, which can vary due to the frequent rearrangements occurring around the *ori-ter* axis through rRNA operon recombination^31^. In our working model, *Vibrio cholerae* O1 biovar El Tor str. N16961, the locus is in the right replichore about 210 Kbp away from *ori1* associated with genes encoding components of the translation machinery (Figure 1). Overall, these observations indicate that the *rplKAJL-rpoBC* locus structure is widely conserved and suggest that, at least among *Vibronaceae*, its genomic position has been selectively maintained throughout the evolution of the clade.

**Figure 1.**
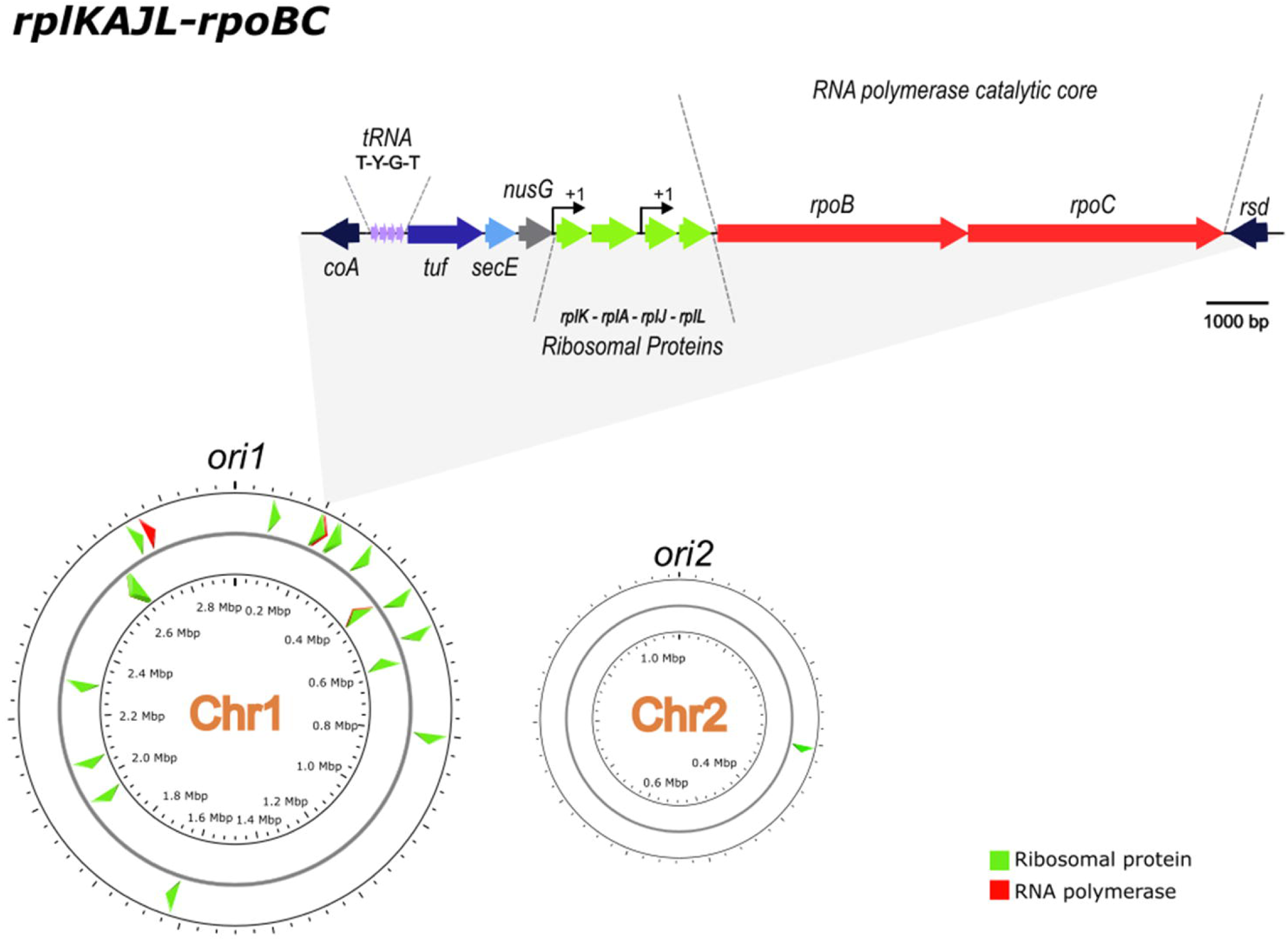
Positional bias of genes involved in the flow of genetic information in *V. cholerae*. The RNA polymerase (red) and ribosomal protein genes (green) tend to be located close to *ori1*. The *rpIKAJL-rpoBC* locus is within the right replichore of the main chromosome. The zoomed section shows the order of the genes (indicated with arrows) within the locus and the promoter regions transcribing *rpoB* and *rpoC*.

### The genome relocation of *rplKAJL-rpoBC* impacts cell physiology in fast-growing conditions

In *V. cholerae, rpoB* and *rpoC* (VC0328-VC0329) are physically linked to the translation-related genes *tuf-secE-nusG-rplKAJL* (VC321-VC0327) and 4 tRNAs (Figures 1 and S1). We relocated the whole locus to several positions along the chromosome to study the impact on cell physiology. The used strategy is despicted in Methods section and Figure S2 performed accordingly to our previous work^14^. This strategy allowed us to move the *rplKAJL-rpoBC* locus to a nearby location, 35 Kbp away from its original position, as well as to the termini of Chr1 (*ter1*) and Chr2 (*ter2*).

We refer to these strains as “movants”^13^ (*i.e.,* isogenic strains where the genomic position of *rplKAJL-rpoBC* changes): Tnp+35, Tnp+1120 and Tnp+479II respectively (Figure 2A). Overall, all strains displayed normal colony morphology and cell shape.

**Figure 2.**
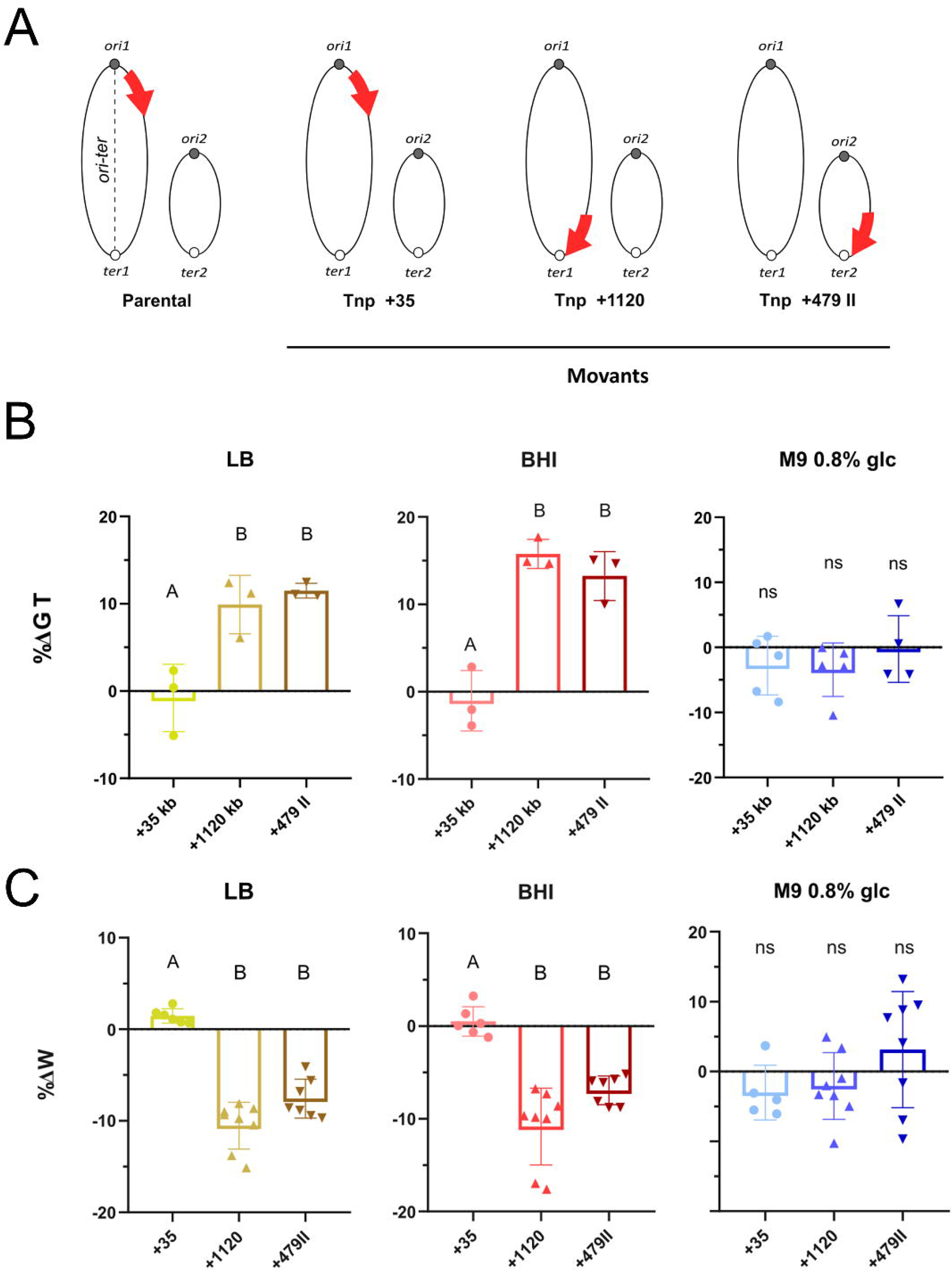
Relocation of *rplKAJL-rpoBC* leads to a nutrient-dependent fitness loss. (A) Genome organization of the movants built in this study. The genomic position of *rplKAJL-rpoBC* is represented by a red arrow. *ori1* and *ori2* are depicted as black circles, while *ter1* and *ter 2* are shown as white circles. (B) The effect of the *rplKAJL-rpoBC* relocation on growth rate (GR) was quantified by averaging the generation time (GT) obtained from at least 3 independent experiments for each movant and normalizing it to the GT of its corresponding parental strain. Results are expressed as percentage of variation (%ΔGT) with 95% CI with respect to parental strains. (C) The fitness (W) of each movant strain was measured by averaging 6 independent experiments and normalizing it to the W of the corresponding parental strain. Results are expressed as percentage of variation (%ΔW with 95% CI with respect to the parental strains. For B and C, statistical significance was analyzed using a one-way ANOVA test. The Holm-Sidak test was then used to compare the mean values obtained for each strain. Letters indicate statistically different (p<0.05) groups.

Fast-growing bacteria tend to cluster their transcription and translation genes near *oriC* possibly for efficient gene expression and replication coordination^8,22^. Based on this, we speculated that the growth rate might be altered in some of the *rplKAJL-rpoBC* movants. To test this, we performed automated growth curves of the parental and the movant strains in different media (Figure 2B, Table S3). In lysogeny broth (LB), the strain Tnp+35 showed no significant changes in its growth while movants in which *rplKAJL-rpoBC* was relocated further from *ori1* exhibited a significant reduction in growth rate of 9.9 ± 1.92 % for Tnp+1120 and 11.5 ± 0.48 % for Tnp+479II. We then tested the strains in a high-nutrient medium that maximizes growth, such as Brain Heart Infusion (BHI). In this case, Tnp+35 growth did not vary significantly from the parental strain. Tnp+1120 and Tnp+479II displayed a slower growth as reflected by a 15.76 ± 0.96 % and 13.25 ± 1.65 % increase of doubling time (Figure 2B). Finally, we tested the growth of movant strains in slow growth conditions using minimal medium (M9 supplemented with 0.8% of glucose). None of them showed growth differences compared to the parental strains.

Overall, the absence of phenotype in the Tnp+35 movant indicates that the precise location of the *rplKAJL-rpoBC* locus does not affect cell physiology and that the relocation process itself is innocuous. However, under fast-growing conditions, relocating the locus far from *ori1* resulted in a growth rate reduction. This phenotype was more pronounced in BHI than in LB, consistent with the latter being a lower-nutrient medium^32,33^. In contrast, no phenotype was observed in minimal medium, indicating that the physiological impact of *rplKAJL-rpoBC* relocation is nutrient-dependent.

Growth curves estimate bacterial population size indirectly through optical density, which cannot distinguish between a reduced growth rate, loss in viability, or changes in light dispersion (e.g., due to altered cell morphology). To assess viability, we plated exponentially growing cells and found no loss, as all strains exhibited a 0.5 × 10□ CFU/OD□□□□□ ratio. We then measured cell size by microscopy in cells grown. In BHI, the length distribution of Tnp+1120 and Tnp+479II was slightly but significantly reduced compared to the parental strain (Figure S3). It is unlikely that this small change in size distribution significantly alters the light dispersion by turbidometry. This observation goes in line with bacterial “growth laws”^34,35^ established decades ago which link cell size to growth rate. Finally, video microscopy of the parental strain, Tnp+35, Tnp+1120, and Tnp+479II confirmed normal cell cycle progression, with no aberrant division or cell death events (Supplementary Videos S1–S4). Meanwhile, the reduction in growth rate became evident.

### Beyond physiology: impact of relocation of *rplKAJL-rpoBC* on fitness

Growth curves provide a broad estimate of cellular fitness. However, they do not account for fitness differences beyond balanced growth and must be interpreted with caution^36,37^. Additionally, they are less sensitive than pairwise competitions^14,15^. To uncover potential fitness differences undetected by growth curves, particularly in minimal medium, we performed pairwise competition experiments. For this, we mixed each movant 1:1 with a *gfpmut3*\*-tagged^38^ *V. cholerae* strain (Table S1). Cultures were diluted and cultured overnight as indicated in the Methods section. Flow cytometry was then used to determine the Gfp^+^/Gfp^-^ ratio before and after overnight pair-wise competition. The number of cells and GFP^+^ ratio at each time point was used to determine absolute fitness (W). W was calculated from these data, and relative fitness (W_rel_), the ratio of each movant’s W to that of its parental strain, was used to compare all movants. We performed the pairwise competition in the three media employed before. The results are summarized in Figure 2C, Table 1 and Table S4. In rich media (i.e. BHI and LB) the Tnp+35 presented a similar capacity to compete against the *gfpmut3*\*-tagged strain, indicating that neither the precise location nor the relocation process impacted fitness. In parallel, the movants where *rplKAJL-rpoBC* locus is relocated far from *ori1*, Tnp+1120 and Tnp+479II, displayed an ∼10% fitness loss (Figure 2C and Table 1). Finally, in minimal medium none of the movants displayed fitness defects (Figure 2C) indicating that the genomic location of *rplKAJL-rpoBC* impacts cell fitness only under fast-growing conditions, closely matching our growth rate observations. This suggests that the primary effect of *rplKAJL-rpoBC* relocation on fitness stems from its impact on the maximum growth rate, while effects on stationary and lag phases appear marginal or negligible.

**Table 1:** Relative fitness (W_rel_) variation of movants under different nutrient abundances. Fitness of *V. cholerae* derivatives were measured by pairwise competition in Lysogeny Broth (LB), Brain Heart Infusion (BHI) and minimal medium M9 supplemented with glucose (M9). Results are shown as mean ± SD with respect to the parental strain.

|  | LB | BHI | M9 1%<br>GLUCOSE |
| --- | --- | --- | --- |
| *rpoBC tnp+35* | 1.014 ± 0.010 | 0.991 ± 0.025 | 0.967 ± 0.039 |
| *rpoBC tnp+1120* | 0.897 ± 0.0260 | 0.895 ± 0.040 | 0.984 ± 0.044 |
| *rpoBC tnp+479 II* | 0.934 ± 0.049 | 0.942 ± 0.028 | 0.967 ± 0.086 |

**Table 2:** Relative fitness variation of merodiploid and Δ*rpoBC* strains in optimal growth conditions. Fitness of *V. cholerae* derivatives measured by pairwise competition against the parental strain carrying *gfpmut3\**. Results are shown as mean ± 95% CI.

| STRAIN | GROWTH CONDITION |
| --- | --- |
|  | BHI |
| **Md($\Delta rpoBC$;+35)** | 0.981 ± 0.020 |
| **Md($\Delta rpoBC$;+1120)** | 0.931 ± 0.038 |

### Genome-wide *rplKAJL-rpoBC* copy-number changes according to its chromosomal location

Despite having the same genetic content, changes in the position of the *rplKAJL-rpoBC* locus affected cell fitness depending on the growth conditions (Figure 2B–C), with the phenotype being more pronounced under conditions that promoted higher replication rates (Table S3).

An increasing number of studies indicate that the genomic location of genes impacts its expression due to replication-associated dosage^20^. These effects arise because bacterial chromosomes replicate from a single *oriC*. Given that the replisome moves at approximately 800 nts/sec^39,40^, genome replication takes about 31 minutes in *V. cholerae.* This is twice as long as its GT of 16 min. This discrepancy can be explained by the phenomenon of multifork replication (MFR), where DNA replication initiates before the completion of the preceding round, leading to the inheritance of partially replicated chromosomes. This ensures that DNA replication maintains pace with population dynamics, thereby preventing the emergence of anucleated daughter cells. Consequently, a decreasing dosage gradient is established along the *ori*-*ter* axis, influencing gene expression. However, there is still some debate around how genomic location alters expression. Some studies point out that expression strength does not correlate well with genomic position within the *ori-ter* axis^13,41,42^. Additionally, subcellular gene positioning may spatially organize the transcriptome, potentially impacting cellular physiology. The fact that relocating *rplKAJL-rpoBC* far from *ori1* affects growth only under fast-growing conditions suggests that replication-linked gene dosage differences may be the mechanism underlying the observed phenotypes^43–45^.

To test this hypothesis, we performed marker frequency analysis (MFA) on the movant strain set (Figure 3A). For this, we cultured the parental and the movant strains in BHI to early exponential phase (OD_600nm_=0.2), after which genomic DNA was extracted and subjected to deep sequencing. We then aligned the reads to the *V. cholerae* reference genome sequence and normalized the coverage to *ter1* since each cell contains a single *ter1* region. As in previous studies^31^, the *oriC* show a maximum coverage and then the Log_2_ number of reads at each locus forms a linear trend between *oriC* and the *ter* in both chromosomes. Changes in coverage can be used to quantify the dosage of every locus at population level during fast growth. The ori1/ter1 ratio provides the maximum dosage difference that can be attributed to genomic location.

**Figure 3.**
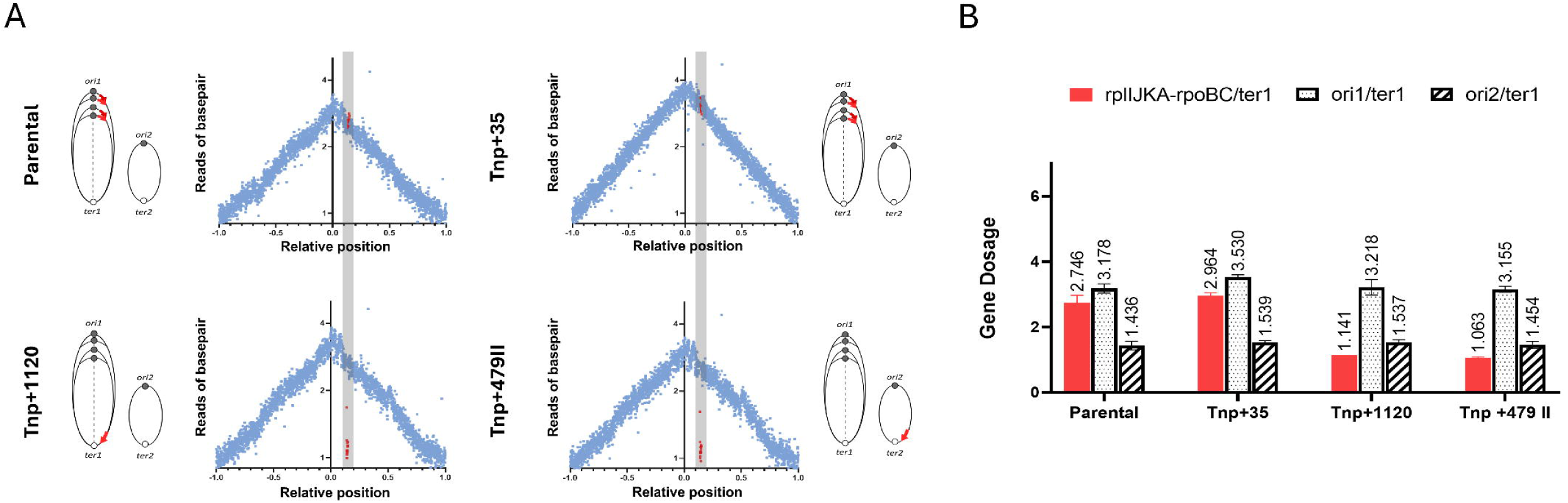
Relocation of *rplKAJL-rpoBC* leads to a genome-wide copy number reduction when this locus is moved to regions near *ter*. (A) Marker Frequency Analysis (MFA) of exponentially growing strains using a corrected reference sequence of Chr1. The log2 of the number of reads starting at each base is plotted against their relative position on Chr1. The position of *ori1* is set to 0 to better represent bidirectional replication. Dots indicate the average of 1,000 bp windows; red dots correspond to *rpIKAJL-rpoBC* genes. Ovals represent the two *V. cholerae* chromosomes and red arrows indicate the *rpIKAJLrpoBC* locus position. (B) Quantification of genome-wide dosage of key loci in the movant strain set. The number of loci per cell of *rpIKAJL-rpoBC* (red), *ori1* (dot pattern) and *ori2* (stripe pattern) was obtained by normalizing the reads to *ter1* coverage in early exponential phase.

In the parental strain grown in BHI, we found 2.746 ± 0.220 copies of the *rplKAJL-rpoBC* locus per cell (*rplKAJL-rpoBC/ter1)*. In the movant Tnp+35, the *rplKAJL-rpoBC/ter1* dosage was 2.964 ± 0.087, which was not significantly different from the parental strain. In contrast, the movants Tnp+1120 and TnpII+479 displayed a single *rplKAJL-rpoBC* copy per cell, with *rplKAJL-rpoBC/ter1* ratios of 1.141 ± 0.001 and 1.063 ± 0.024, respectively. The ori1/ter1, ori2/ter1 ratios and genome coverage of any of the movants did not significantly change from the parental strain (Figure 3A, B). These findings indicate that relocating *rplKAJL-rpoBC* far from *ori1* reduces the genome-wide copy number of the locus but does not affect the DNA replication process.

### RNA polymerase expression changes according to *rplKAJL-rpoBC* genomic position

Next, we assessed whether the genome-wide copy number reduction of *rplKAJL-rpoBC* affected RNAP expression. The *rplKAJL-rpoBC* locus consists of two ribosomal protein operons, *rplJ* and *rplK,* followed by *rpoB* and *rpoC* which encode the RNAP core (Figure 1). These latter are under the transcriptional control of the ribosomal protein genes. To quantify RNAP expression we tagged RpoC (β’ subunit) by building a *rpoC*-mCherry translational fusion in the parental and in the Tnp+1120 movant.

Tagging the RNAP β’ subunit did not affect the previously observed phenotypes, as the Tnp+1120-mCherry movant displays a significantly slower growth than the parental *rpoC-mCherry* strain (Figure S4). Then, the labeled strains were subjected to flow cytometry to quantify the intensity of mCherry which correlates with the abundance of the RpoC. As shown in Figure 4A, relocation of *rplKAJL-rpoBC* to *ter1* resulted in a –22.84 ± 5.400% reduction in mCherry signal under optimal growth conditions and a –6.242 ± 2.600% reduction under slow-growth conditions (M9 with 0.8% glucose).

**Figure 4.**
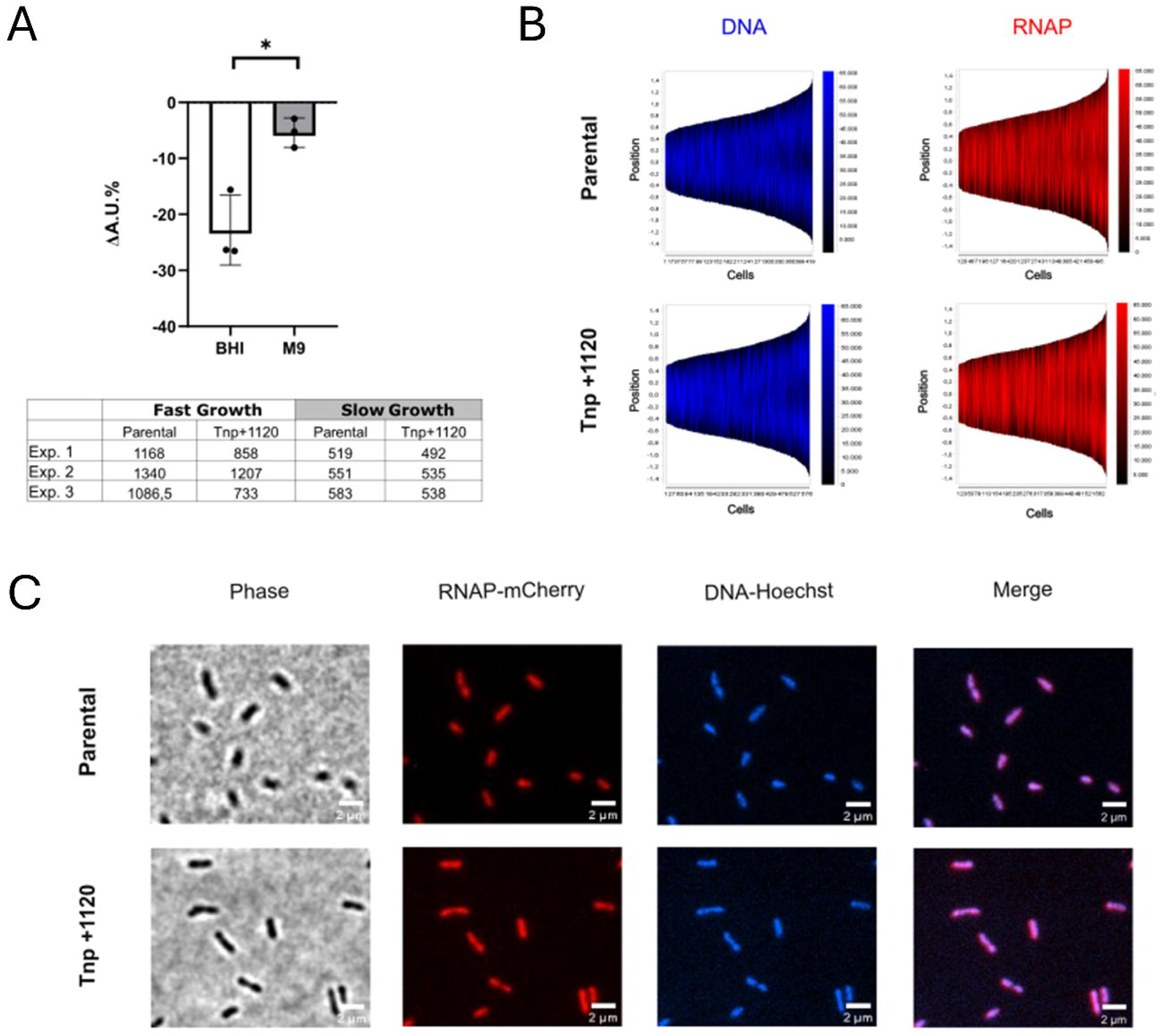
Distribution of RNAP in *rpIKAJL-rpoBC* movant. **(A)** Difference in arbitrary fluorescence units (A.U.) of strain Tnp+1120-mCherry compared to Parental-mCherry from 3 different experiments under fast (BHI) and slow (M9 0.8% glucose) growth conditions. Statistical significance was analysed by two-tailed t-test (p>0.05; *, p<0.05; **, p<0.01; ***, p<0.001;****, p<0.0001). The table at the bottom shows the absolute values of A.U.of a representative experiment. **(B)** Demographic representation of DNA and RNAP signals in parental and Tnp+1120 strains made with MicrobeJ image analysis. 400-500 cells were sorted by size (from smallest on the left to largest on the right). Fluorescence intensities are shown as a heatmap with minimum and maximum values shown on the vertical scale. **(C)** Spatial distribution of *rpoC* fused to *mCherry* with respect to the nucleoid of *V. cholerae* parental and Tnp+1120 cells: phase-contrast images in the first panel; RNA polymerase RpoC subunit tagged with mCherry in the second panel; nucleoid signal was obtained by staining DNA with Hoescht in the third panel. The RNAP-red and DNA-blue channels were merged (fourth panel) to identify the relative positions of RNAP relative to the nucleoid. Scale bar, 2 mm.

### Phenotypes associated with *rplKAJL-rpoBC* genomic position are driven by gene dosage changes

In the previous sections, we showed that relocating *rplKAJL-rpoBC* away from *ori1* reduces its dosage during exponential phase in rich medium and decreases total RNAP levels. Thus, the most plausible explanation for the observed changes in growth rate and fitness is the dosage effect associated with MFR at each chromosomal position. However, alternative mechanisms could also contribute to the observed phenotype. For instance, *rplKAJL-rpoBC* relocation may alter the subcellular positioning of its gene products, potentially impacting their interactions with other factors required for macromolecular complex formation, such as ribosome assembly (*rplJ* and *rplK*) or RNAP assembly and promoter binding (*rpoB* and *rpoC)*^44^. Additionally, heterologous positioning of *rplKAJL-rpoBC* might be intrinsically detrimental if it disrupts the expression of neighboring genes. Finally, inserting highly expressed genes at new chromosomal locations could locally affect chromosome conformation^46^.

To determine whether *rplKAJL-rpoBC* relocation affects the subcellular distribution of RNAP, we grew the parental and Tnp+1120 strains expressing β’-mCherry in BHI until early exponential phase. After fixing the cells, we stained the DNA with HOESCH 33342 and imaged them using Differential Interference Contrast (DIC) and epifluorescence microscopy, recording the channels corresponding to the DNA and the RNAP signals (Figure 4B, C). Since γ-Proteobacteria such as *V. cholerae* divide by binary fission, cell length correlates with cell age. Accordingly, demographs depicting DNA and RNAP signals as a function of cell length provide an approximate representation of the cell cycle (Figure 4B). Cells exhibited a strong DNA signal at the center, corresponding to the nucleoid, while longer cells displayed two segregating DNA foci prior to division. The mCherry signal closely followed the DNA signal within the cellular space throughout the cell cycle (Figure 4B and S5). As both strains exhibited similar RNAP localization patterns, these results strongly suggest that *rplKAJL-rpoBC* genomic positioning does not affect the subcellular distribution of RNAP in *V. cholerae*.

To separate the effects of chromosomal position from genome-wide *rplKAJL-rpoBC* copy number, we constructed merodiploid strains. For this, we introduced a second copy of the locus through natural competence-mediated transformation, employing gDNA from a strain bearing *rplKAJL-rpoBC* at an alternative location (described in Methods section). To ensure that the observed phenotypes were not due to the accumulation of suppressor mutations, we independently generated these strains multiple times. First, we constructed a strain carrying *rplKAJL-rpoBC* at its original location and a second copy 35 Kbp downstream, designated Md(0;+35). This strain allowed us to test the effect of artificially increasing locus dosage, as both copies remained close to *ori1* (Figure 5A). Next, we built Md(+35;+1120), a strain with one copy at position +35 and a second copy relocated far from *ori1* at position +1120. This merodiploid strain allowed testing whether the relocation of *rplKAJL-rpoBC* exerts a dominant effect over the copy at its original location. Finally, we constructed Md(+1120;+479II), a strain bearing two *rplKAJL-rpoBC* copies far from *ori1,* at the termini of both chromosomes. This merodiploid enabled us to determine if the impact of *rplKAJL-rpoBC* relocation far from *ori1* could be compensated solely by an increase in dosage, independent of chromosomal location. These strains were evaluated through automated growth curves under fast-growing conditions. As shown in Figure 5 B and C, the Md(0;+35) strain showed no alteration in growth rate compared to the parental or Tnp+35 strain, despite the increased *rplKAJL-rpoBC* dosage. We interpret that, at least in these conditions, an excess of *rplKAJL-rpoBC* dosage is not detrimental to cell physiology and does not limit growth. When we tested Md(+35;+1120), we observed an increased growth rate compared to the movants with *rplKAJL-rpoBC* relocated far from *ori1*. This indicates that *rplKAJL-rpoBC* at its original location has a dominant effect over the copy positioned at the chromosomal termini, suggesting that the physiological impact of *rplKAJL-rpoBC* relocation is due to dosage effects rather than an intrinsic disadvantage of the heterologous locus position. Interestingly, Md(+1120;+479II) exhibited a generation time similar to that of the parental strain and a statistically significant increase in growth rate compared to the Tnp+1120 and TnpII+479 movants. Overall, these experiments demonstrate that the phenotypes observed upon *rplKAJL-rpoBC* locus relocation result primarily from genome-wide copy number variations, which prevail over potential dosage-independent effects.

**Figure 5.**
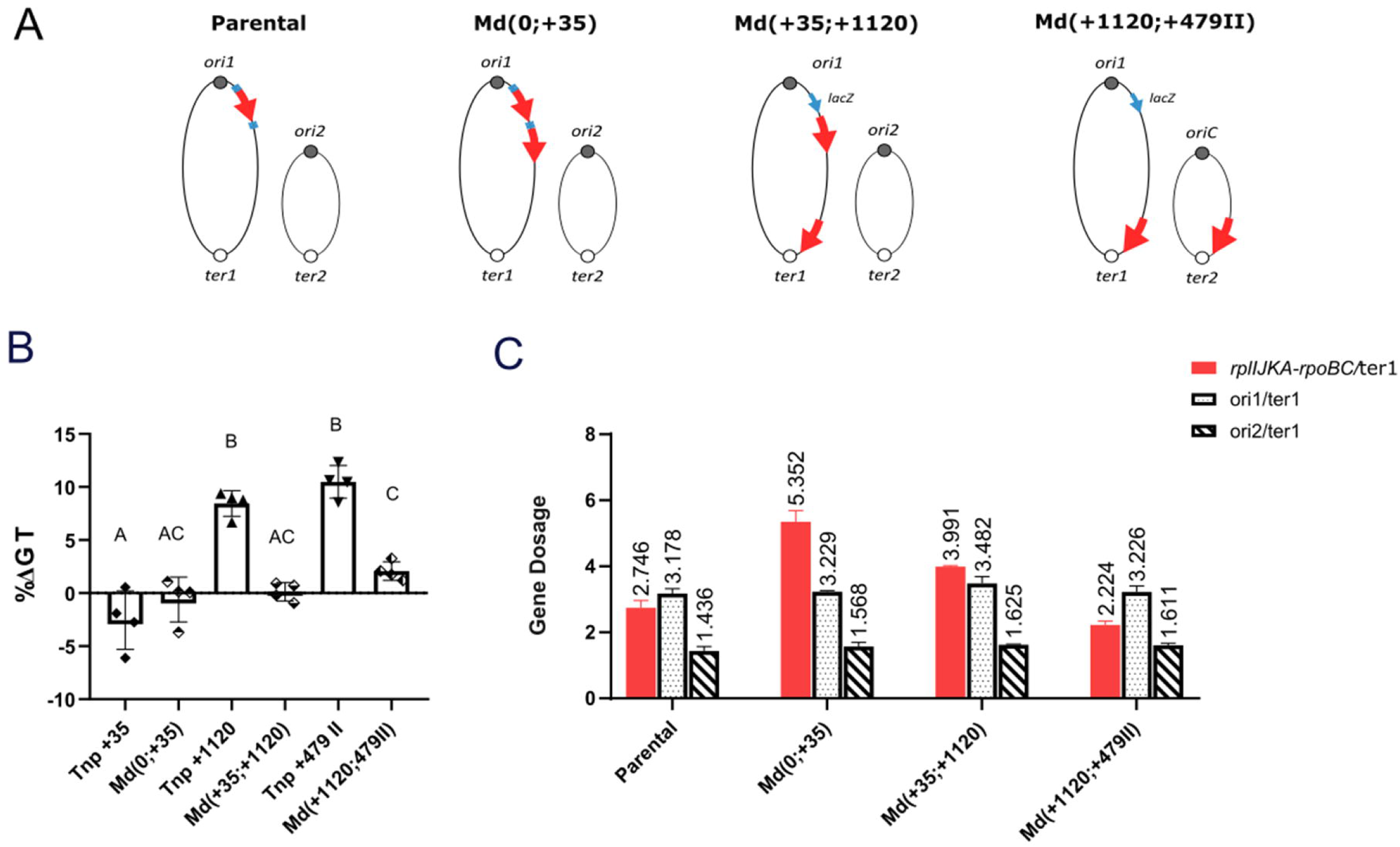
The addition of a second copy of the *rpIKAJL-rpoBC* locus restores growth regardless of chromosomal position. (A) Scheme of generated merodiploid strains. Ovals represent chromosomes, red and blue arrows represent the *rpIKAJL-rpoBC* locus and the selection marker, respectively. (B) The GR of merodiploid parental was quantified by averaging the GT obtained in at least 4 independent experiments, normalizing it to the GT of the corresponding parental strain. Results are expressed as percentage of variation (%ΔGT) with 95% CI with respect to the parental strains. Statistical significance was analyzed using a one-way ANOVA test. The Holm-Sidak test was then used to compare the mean values obtained for each strain. Letters indicate statistically different groups. (C) Number of reads of *rpIKAJL-rpoBC* (red), *ori1* (dot pattern), and *ori2* (stripe pattern) normalized to *ter1* in early exponential phase movant strains obtained in 2 independent experiments.

**Figure 5.** The addition of a second copy of the locus restores growth in strains regardless of chromosomal position. (A) Scheme of generated merodiploid strains. Ovals represent chromosomes, red arrows represent *rpIKAJL-rpoBC* locus and the selection marker in light blue. (B) The GR of merodiploid parental was quantified by averaging the GT obtained in at least 4 independent experiments normalizing it to the GT of the corresponding parental strain. Results are expressed as percentage of variation (%ΔGT) with 95% CI with respect to the parental strains. Statistical significance was analysed using a one-way ANOVA test. The Holm-Sidak test was then used to compare the mean values obtained for each strain. Letters indicate statistically different groups. (C) Number of reads of *rpIKAJL-rpoBC* (red), *ori1* (dot pattern) and *ori2* (stripe pattern) normalized to *ter1* in early exponential phase movant strains obtained in 2 independent experiments.

### The change in dosage of RNA polymerase genes within the *rplKAJL-rpoBC* locus explains the phenotypes observed

The main hypothesis of our work is that the conserved genomic location of RNAP provides a fitness advantage that has been selected during the evolution of the clade. However, beyond the RNAP core, the *rplKAJL-rpoBC* locus also encodes genes involved in the translation process, such as RPs, tRNAs, and *nusG*. To determine whether the observed phenotypes arise specifically from the RNAP core genes or from other genes within the locus, we employed a genetic approach using the Md(0;+35) and Md(0;+1120) strains, considering that *rpoB* and *rpoC* are located at the 3’ end of the locus (Figure 6A). In each merodiploid strain, we deleted the *rpoB* and *rpoC* genes at the *ori1*-proximal site, generating Md(Δ*rpoBC*; +35) and Md(Δ*rpoBC*; +1120), respectively, using allele replacement (Figure 6A,B and Methods section). As shown by MFA, we successfully uncoupled the dosage of the two gene groups, as Md(Δ*rpoBC*; +1120) exhibits a change exclusively in the dosage of *rpoB* and *rpoC*, while the rest of the locus remains unaffected (Figure 6B-C and Figure S6). If the RNAP genes contribute to the previously observed phenotypes, Md(Δ*rpoBC*; +1120) should display slower growth and a reduced fitness compared to Md(Δ*rpoBC*; +35). Conversely, if the genomic location of the translation-related genes within the *rplKAJL-rpoBC* locus determines *V. cholerae* growth and fitness, both strains should show similar phenotypes. To test this, we used automated growth curves to compare the generation times of the selected merodiploid strains with different Δ*rpoBC* derivatives grown under fast-growing conditions. As shown in Figure 6D, consistent with previous experiments, Tnp+35 exhibits no significant difference in GT compared to the parental strain, whereas Tnp+1120 shows a 10.83 ± 2.55% increase in doubling time. Meanwhile, Md(Δ*rpoBC*; +35) displays a similar growth rate to Md(0;+35) (–2.814 ± 3.655%), while the Md(Δ*rpoBC*; +1120) derivative shows a 6.66 ± 2.81% increase in GT. In parallel, pairwise competition experiments reveal that Tnp+1120 undergoes a 10.84 ± 4.14% fitness loss, while deletion of the *rpoB* and *rpoC* copy proximal to *ori1* results in a –7.33 ± 4.17% fitness loss (Figure 6E). Finally, the Md(Δ*rpoBC*; +35) strain shows a fitness value comparable to the parental strain (–1.97 ± 1.79%).

**Figure 6.**
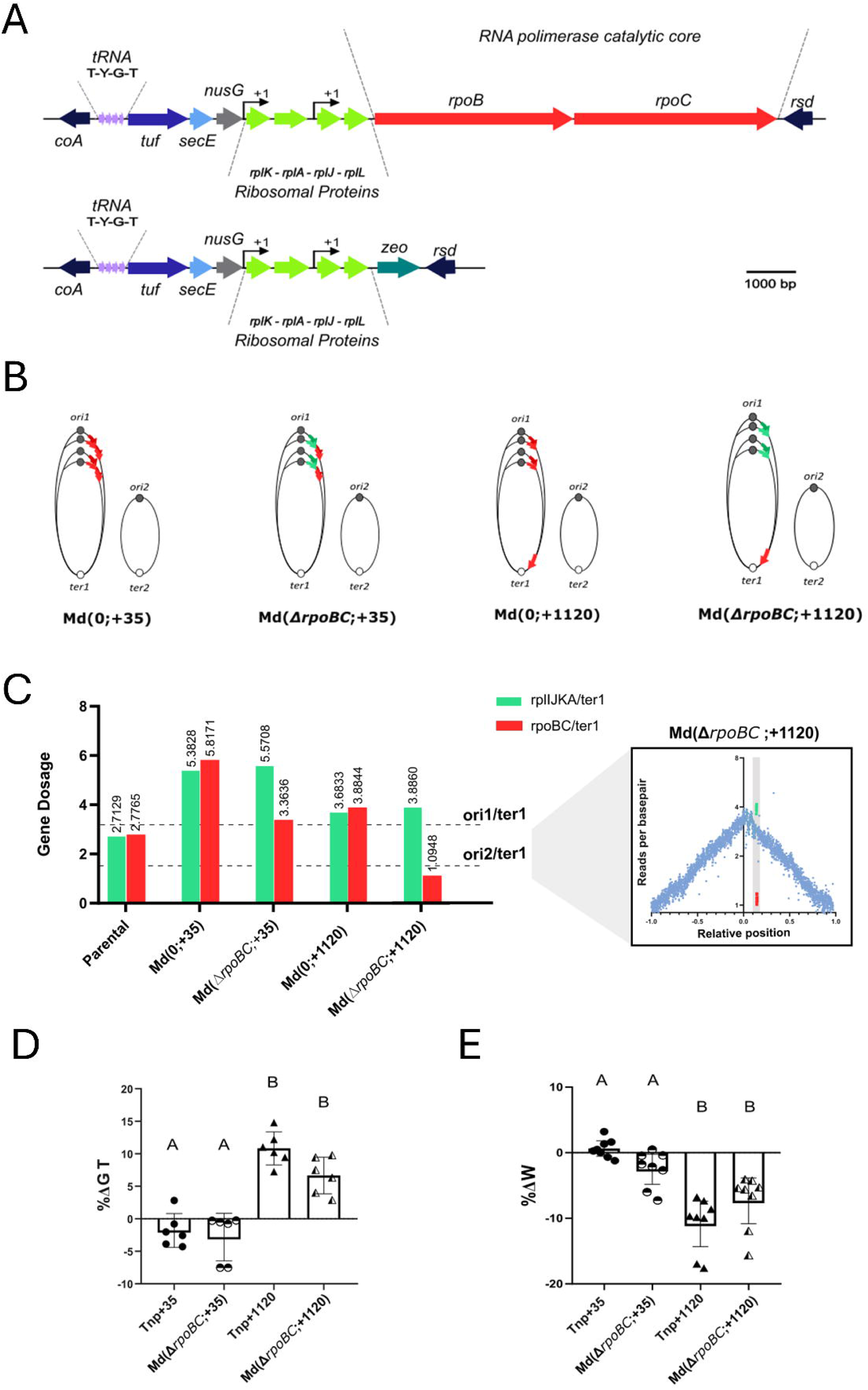
*rpoB* and *rpoC* dosage impacts *V. cholerae* physiology. (A) The scheme shows the genetic structure of wild type *rpIKAJL-rpoBC* (upper panel) and the locus deprived of *rpoB* and *rpoC* by deletion using a Zeo^R^ marker (lower panel). (B) Graphical representation of *V. cholerae* merodiploid strains genomes. The *rpIKAJL-rpoBC* and *rpIKAJL-zeo* are displayed in red and green respectively during MFR. (C) The number of reads of *rpIKAJL* (green) and *rpoBC* (red) genes were normalized to *ter1* in early exponential phase. One representative experiment conducted in BHI medium is shown. The dotted lines represent the average gene dose of each *ori* relative to the *ter1* on each tested strain. On the right, a representative MFA experiment of Md(Δ*rpoBC*;+1120) showing how *rpoB* and *rpoC* dosage is uncoupled from the genome-wide copy number of the rest of the locus. (D) The effect of *rpoB* and *rpoC* relocalization on GT was quantified in automated growth curves. The difference in TG of each movant with respect to its parental strain was calculated in at least 5 independent experiments. Results are expressed as percentage of variation (%ΔGT) with 95% CI with respect to the parental strains. (E) The fitness (W) of each movant was measured by averaging 6 independent experiments and normalizing it with respect to the W of the corresponding parental strain. Results are expressed as percentage of variation (%ΔW) with 95% CI with respect to the parental strains. Statistical significance was analyzed using a one-way ANOVA test. The Holm-Sidak test was then used to compare the mean values obtained for each strain. Letters indicate statistically different groups.

In summary, these results demonstrate that the reduction in *rpoB* and *rpoC* dosage within the *rplKAJL-rpoBC* locus is, at least in part, responsible for the chromosomal positional effects observed.

## Discussion

An increasing number of studies indicate that gene location within the bacterial chromosome is non-random^9,10,20^. The role of chromosomal gene order in cellular homeostasis is particularly relevant for understanding chromosome architecture, cellular function, and the design of synthetic chromosomes^47–49^. More specifically, gene order within the bacterial genome may coordinate key interconnected processes, including chromosome organization, cell cycle progression, DNA replication, and, notably, gene expression regulation.

### Positional genetics: Potential Applications and Limitations

The expression of genetic information is crucial for life and is tightly linked to cell growth. In this regard, RNAP abundance and transcriptional activity is coordinated with cellular size both in prokaryotes^21,50^ and eukaryotes^51,52^. Our experiments closely match these recent observations since most affected *rplKAJL-rpoBC* movants displayed a smaller size (Figure S3). The most affected movants showed a nutrient-dependent growth rate reduction of about 15% (Figure 2). We demonstrate that the observed phenotypes were dosage-dependent (Figures 3 and 5) and, at least in part, influenced by the chromosomal location of RNAP genes (Figure 6). *rpoB* and *rpoC* exert a global control of cell physiology and metabolism ^53^. Izard and colleagues previously engineered an *Escherichia coli* strain in which *rpoB* and *rpoC* expression was tightly regulated by an heterologous promoter^23^. This genetic switch allowed turning on and off cell transcription. When turned off, cell metabolism remained active, allowing to uncouple growth from metabolite production. It was effectively a switch since in different RNAP inductor concentrations, growth rate was either maximal or null. Although this switch was stable, some escape events were detected after 48hs of culture. This contrasts with positional genetics as a strategy to regulate cell growth: *rplKAJL-rpoBC* relocation to chromosome termini led to intermediate phenotypes (i.e. 10-15% growth rate reduction, Figures. 2B and 6D). Also, the genomic location is likely to be more robust. Previous work on relocating the S10 locus showed no escape events, even after 1,000 generations of adaptive laboratory evolution^17^. These complementary approaches highlight the importance of *rpoB* and *rpoC* expression in global regulation and demonstrate that modulating their expression through different strategies can be an effective approach for reprogramming or fine-tuning growth for biotechnological applications. They also expand our understanding of how specific aspects of the main genes of the transcription machinery, such as genomic location, subcellular distribution and levels of expression contribute to cell functioning. This issue is key for understanding the structure-functional relationships of bacterial chromosomes.

The bacterial chromosome is able to tolerate large inversions and other major genomic modifications with little or no physiological impact ^47,54^. It has alsobeen shown that it is possible to impair cellular functions such as growth, induction of natural competence for transformation and sporulation by relocating genes of these processes^11,12,14,55^. However, gaining functions or improving cellular processes seems unlikely. For instance, in this work we added extra copies of the *rplKAJL-rpoBC* locus without a noticeable improvement in cell growth or fitness despite the increase in gene dosage (Figure 5 and S6). This was also observed in cells harboring additional copies of the S10 locus^15^ or the *rrn* operon^56^. A further increase in the dosage of these loci was shown to be detrimental to bacterial physiology^15,56^. We do not rule out the possibility of gaining function through more subtle strategies. For example, it would be worth testing whether relocating the *rplKAJL-rpoBC* locus closer to *oriC* or adding additional copies reduces doubling time in a slow-growing model. Positional genetics and rational genome engineering could be valuable strategies for accelerating cell division in slow-growing bacteria, which may have significant biotechnological applications.

### The genomic location of RPs and RNAP genes influences cell physiology through different mechanisms

Previous work from our group demonstrated that the proximity of ribosomal protein genes to *ori1* optimizes growth, fitness, and infectivity in *V. cholerae*^14^. This trait is highly stable over long-term evolution, as shown by adaptative laboratory evolution^17^. In this paper, we provide a novel example of a gene cluster whose genomic position contributes to cell homeostasis, at least in optimal growth conditions. We studied the core transcription genes because they are upstream the translation genes in the central dogma^57^. Pioneering work from Eduardo Rocha’s Lab indicates that, when predicting doubling time based on the genome sequence, the location of RNAP genes has higher predictive power than the chromosomal location of RP genes^22^. Thus, we expected that relocating RNAP genes far from *ori1* would have a greater impact on cell physiology than moving RP genes. Unexpectedly, our experiments suggest the opposite since 1) When relocated to the chromosomal terminus, the S10 locus exhibits a 15-17% growth deficit, whereas *rplKAJL-rpoBC* relocation results in a 10% increase in doubling time in LB (Figure 2B); 2) S10 movants are less competitive than their parental strains in the absence of MFR, while *rpoBC*+1120 and *rpoBC*+479II are not outcompeted by their parental strains in slow growing conditions (Figure 2C). A possible explanation for this discrepancy is that the physiological impact of relocating the S10 and *rplKAJL-rpoBC* loci far from *ori1* involves different molecular mechanisms. In the case of S10 relocation, the physiological effects were primarily associated with changes in cytoplasmic macromolecular crowding^16^. For example, S10 mutants displayed altered replication dynamics, and their physiological deficits were alleviated under hyperosmotic conditions. In contrast, *rpoBC* movants did not display alterations in replication dynamics (Figures 3 B and 5 C). In line with this, recent modeling of autocatalytic cycles in bacteria shows that a reduction in growth rate due to shortage in RNAP occurs without affecting the ribosomal protein mass fraction^57^. Elucidating the mechanistic differences between RNAP and RP gene relocation requires further investigation, which is beyond the scope of this study. The transcriptomic characterization of the *rpoBC* movants and performing the single or dual relocation of *rplKAJL-rpoBC* and S10 loci within the same genetic background will provide further insight into this issue.

### Gene dosage, spatial address and genetic linkage of transcription-translation genes

We determined that *rplKAJL-rpoBC* locus chromosomal position recruits MFR for adaptative purposes by increasing RNAP gene dosage to increase expression of transcriptional machinery components when most needed, namely during the exponential growth phase(Figures 2, 5 and 6). While we attributed most of the observed effect to the genomic position of RNAP genes, it is intriguing that transcription, translation and protein secretion genes are genetically and physically linked. Notably, the *rplKAJL-rpoBC* and S10 loci encode and share functionally related genes involved in secretion (*sec*), transcription (*rpo*), translation (*rpl/rps*), and transcription-translation coupling (*nus*). Recent studies have shown that RNAP and the ribosome can form a single complex known as the ‘expressome’^58,59^. The S10 and *rplKAJL-rpoBC* loci exemplify how operon structure facilitates the physical proximity required for the assembly of protein multimers while transcription and translation are still in progress, as demonstrated in the case of luciferase^59^. This could explain why *rplKAJL-rpoBC* and S10 are highly conserved across bacteria, archaea, and certain eukaryotes. Studies from other laboratories have explored the importance of subcellular localization of RNAP^60^ and RP^61^. We did not find major alterations in RNAP distribution (Figure 4) but we do not rule out the possibility that more subtle effects can be unraveled using super-resolution microscopy and cellular microbiology techniques for studying the subcellular location of loci such as S10 or *rplKAJL-rpoBC*, their transcripts, and gene products. These approaches will provide insights into the spatial organization of the central dogma, a topic that has only recently begun to be explored^62^. However, for transcription and translation genes, we believe that their high cellular concentration may hide the impact of subcellular localization. Indeed, our cell biology experiments showed that RNAP distribution closely follows the nucleoid, indicating, consistent with previous reports in *E. coli*, that the *rpoC*-mCherry translational fusion is fully functional^23,60^.

The influence of genomic location on the behavior of key genetic loci appears to vary depending on the model employed^63^. Other genetic loci, such as those encoding nucleoid-associated proteins, may be influenced by genomic location independently of genome-wide gene dosage^13^. The impact of differential transcript localization, particularly for essential yet lowly expressed genes, remains to be explored^43–45^. In this vein, we envisage employing microscopy techniques to simultaneously tag and track specific genetic loci, their mRNA, and fluorescently labeled gene products over time (Figure 4). This approach will allow us to test the concept of gene ‘spatial address’ and its potential contribution to cellular function^64^. The ‘spatial address’ of genes may help to localize their encoded proteins to specific subcellular regions, facilitate interactions with functional partners, or create localized microconcentrations within the highly crowded bacterial cytoplasm.

### The link between *rplKAJL-rpoBC* genomic location and the ecophysiology of *V. cholerae*

Rapid growth in bacteria is associated with specific genomic signatures: a high number of rRNA operons and tRNAs, optimized codon usage patterns, and the clustering of genes involved in genome expression near *oriC*^65^. Conversely, distinct genome organization patterns are linked to slow growth^66^. The microbial growth regime, either fast or slow, is linked to the environment where microbes live. The availability of nutrients may vary widely in the environment. The microorganism adapted to low concentration of nutrients are called oligotrophs in contrast to copiotrophs that thrive in high nutrient concentrations. Copiotrophs and oligotrophs are associated with fast and slow growth respectively^67^. Oligotrophs typically exhibit slow growth, maximizing resource efficiency, whereas copiotrophs grow rapidly, enabling them to exploit transient periods of high nutrient availability. Since environmental nutrient levels fluctuate, copiotrophic microbes often experience feast-and-famine cycles. During nutrient surges, rapid growth allows them to outcompete other organisms, while oligotrophs persist by efficiently utilizing scarce resources^68^.

The life cycle of *V. cholerae* includes a host-associated phase within the gut, where it is exposed to high nutrient concentrations, and a free-living phase in aquatic environments with limited organic matter^69^. Its genomic organization aligns with that of a fast-growing bacterium (copiotrophic). In this study, we experimentally validated this bioinformatically predicted correlation. Although the observed effects were smaller than expected, relocating a 14 Kbp region encoding *rplKAJL-rpoBC* led to significant physiological changes within this isogenic strain set (Figures 1 and 2). We demonstrated that, in fast growing conditions the *rpoB* and *rpoC* chromosomal location was important for achieving maximal growth and fitness. Since the chromosomal position of these genes is conserved in all *Vibrionaceae* genomes (Table S2), it is likely a shared adaptive trait within the clade. The observed effects were nutrient-dependent: strains with *rplKAJL-rpoBC* relocated near the chromosome terminus exhibited a more pronounced growth defect in BHI than in LB, relative to the parental strain. However, we did not detect significant physiological effects in minimal medium (Figures 2B and 2C). Nevertheless, we cannot rule out a role for *rplKAJL-rpoBC* chromosomal location during famine periods since we measured fitness in minimal medium in pairwise competitions performed for relatively few generations. Notably, *rplKAJL-rpoBC* relocation away from *ori1* resulted in ∼5% lower RNAP levels (Figure 5C). A subtle but persistent fitness effect over many generations could provide an adaptive advantage, which we did not assess in this study. Here, we focused on growth rate and fitness without evaluating carbon efficiency (*i.e.*, the number of carbon moles utilized per newly generated cell). While this parameter has been extensively studied in the context of *rrn* ploidy, it remains an open question in our system.

Altogether, our results suggest that the genomic location of RNAP genes provides an advantage during periods of high nutrient availability, facilitating rapid niche colonization. Thus, the chromosomal positioning of *rplKAJL-rpoBC* may represent an adaptation to the life cycle and ecology of *vibrios*, optimizing growth in host-associated environments where organic matter is abundant and rapid colonization is crucial. Testing the performance of *rplKAJL-rpoBC* movants in an animal model would be a valuable avenue for future research.

## Conclusions

In this study, we demonstrated that the genomic location of the *rplKAJL-rpoBC* locus influences cell physiology and fitness. Growth impairment caused by ectopic relocation of *rplKAJL-rpoBC* highlights the importance of gene positioning in bacterial chromosome organization. Our findings emphasize the role of spatial genome architecture in cellular function and adaptation. Understanding these principles will contribute to the study of complex biological systems and have significant implications for genome design, bioengineering, and biotechnology.

## Methods

### Bacterial strains and plasmids

All strains derive from *V. cholerae* serotype O1 biotype El Tor strain N16961 with is Str^R^^70^. Strains and plasmids used in this study are listed in Table S1.

### Culture conditions

Bacterial cultures were cultured in Lysogeny Broth (LB, Oxoid), unless otherwise specified, supplemented with appropriate antibiotics. Streptomycin (Str, 100 μg/mL), chloramphenicol (Cm, 3 μg/mL), kanamycin (Km, 25 μg/mL), spectinomycin (Spec, 100 μg/mL), carbenicillin (Carb, 50 μg/mL), Trimethoprim (Trim,5 μg/mL) and zeocin (Zeo, 25 μg/mL) were added when required. For fast growing conditions we employed LB or Brain Heart Infusion (BHI, Britannia, Argentina), at 37°C agitating at 200-250 rpm. M9 medium supplemented with 0.8% glucose, CaCl_2_ (5 mM), MgSO_4_ (32 mM), thiamine (4 μM) and micronutrient solution (EDTA 13.4 mM, FeCl_3_-6H_2_O 3.1 mM, ZnCl_2_ 0.62 mM, CuCl_2_-2H_2_O 76 µM, CoCl_2_-2H_2_O 42 µM, H_3_BO_3_ 162 µM, MnCl_2_-4H_2_O 8.1 µM.) was used for slow-growing conditions.

### General procedures

Genomic DNA (gDNA) was extracted using the GeneJET Genomic DNA Purification Kit while plasmid DNA was extracted using the GeneJET Plasmid Miniprep Kit (Thermo Scientific) or ADN Puriprep B and P (InbioHighway, Argentina). PCR assays were performed using DreamTaq, Phusion High-Fidelity PCR Master Mix (Thermo Scientific) or MINT 2X (InbioHighway, Argentina). Oligonucleotides were purchased from Macrogen, Ko.

### *rplKA-JL-rpoBC* movant strain generation

The *rplKA-JL-rpoBC* locus in *V. cholerae* spans from coding sequences (CDs) VC322 to VC329. The *attL*_λ_ site was introduced between VC319 and VC320 and *attR*_λ_ site between VC330 and VC331 using natural transformation^71^. The insertion of recombination sites was made as in previous study^14^. Briefly, *attL*λ and *attR*λ linked to Km^R^ and Cm^R^ were amplified from plasmids pMP98 and pMP102 respectively. These products were linked to amplicons of ∼1Kbp upstream and downstream to their insertion site using PCR assembly (Table S5 and Figure S2). The VG0052 corresponds to the *V. cholerae* strain where *rplKAJL-rpoBC* locus is flanked by *attL*_λ_ and *attR*_λ_. A 33bp-long *attB*’ site linked to Trim^R^ was amplified from pASB9 (Table S5). Then it was joined to upstream and downstream homology regions by PCR assembly for each target region. In all cases, *attB*’ sites were inserted at intergenic zones of inversely oriented ORFs to avoid polar effects. Parental strains bearing *attL*_λ_, *attR*_λ_ and *attB*’ (Table S1) were transformed with plasmid pTSA29CXI^72^ and incubated in LB at 37°C until reaching an OD_600nm_ of 0.3-0.5 to promote the expression of Xis_l_ and Int_l._ The excisive recombination *attL*_λ_×*attR*_λ_reconstitutes *lacZ*. Selection was performed by plating on LB-agar supplemented with 5-Bromo-4-chloro-3-indolyl β-D-galactopyranoside (X-gal, 100μg/mL) or M9 with 1% lactose. Candidates were confirmed as *rpIKA-JL-rpoBC* movants if they met the following criteria i) blue colony or Lac^+^ phenotype ii) PCR confirmation of *lacZ* amplification, iii) PCR confitmation of *attB*’ absence and iv) PCR confirmation of genes adjacent to the relocated *rpIKA-JL-rpoBC* locus.To ensure that possible phenotypes were not caused by suppression or fortuitous mutations occurring in single clones during the construction process, we generated these movants several times independently.

### *rpIKA-JL-rpoBC* merodiploids and *ΔrpoBC* strain generation

Parental and *rplKA-JL-rpoBC* positional mutants were transformed with pMP109 plasmid (Table S1) and selected on LB agar plates supplemented with carbenicillin. Transformed clones were cultured at 37°C until reaching an OD_600nm_ of 0.3-0.5 and then supplemented with 0.5% of arabinose overnight (ON) to induce *aph* and *cat* removal via recombination reaction catalyzed by Flp and Cre recombinases. Individual clones were then selected from LB-agar plates and restreaked on LB plates supplemented with kanamycin, chloramphenicol and streptomycin. Clones showing a Kan^S^ Cm^S^ phenotype were subsequently grown ON in liquid LB at 37°C to promote pMP109 plasmid loss. Cultures were restreaked on LB-agar plates to obtain isolated clones. Selected strains were induced for natural competence^71^ and transformed with gDNA from Tnp+35 for Md(0;+35), with gDNA from Tnp+1120 for Md(0;+1120) and Md(+35;+1120), or gDNA from Tnp+479II for Md(0;+479 II) and Md(+1120;+479 II). Cm^R^ and Kan^R^ clones were selected on LB-agar plates supplemented with the appropriate antibiotics. Genotypes were further confirmed by PCR and whole genome sequencing. To generate *ΔrpoBC* strains, merodiploid strains Md(0;+35) and Md(0;+1120) were transformed with pMP109 to induce the excision of *aph* and *cat* genes as described before. The resulting Kan^S^ Cm^S^ strains were induced for natural compentence and transformed with *rplKAJL*-*zeo*-*vc330* DNA fragment (Figure 6). Cm^R^ and Kan^R^ clones were selected on LB-agar plates supplemented with the appropriate antibiotics. The genotype was further confirmed by PCR.

### *rpoC-mCherry* strain generation

DNA cassettes containing the *mCherry* gene were transferred from a plasmid vector to the chromosome through two homologous recombination steps^73^. Briefly, tworegions spanning the point of insertion were amplified from Parental+1120 Δ(*aph,cat*) and *rpoBC* Tnp+1120 Δ(*aph,cat*) chromosomal DNA by PCR to provide homology for integration. The amplified fragments were cloned into an R6K c-ori-based suicide vector, pMP7, which encodes the *ccdB* toxin gene under the control of an arabinose-inducible and glucose-repressible P_BAD_ promoter^74^. The mCherry fluorescent protein sequence was then cloned between the two chromosomal fragments. For cloning, π3813 was used as a plasmid host and for conjugal transfer of plasmids to *V. cholerae* strains, *E. coli* β3914 was used as the donor^73^. Selection of the plasmid-borne drug marker resulted in integration of the entire plasmid in the chromosome by a single crossover event. Elimination of the plasmid backbone resulting from a second recombination step was selected by arabinose induction of the *ccdB* toxin gene. Colonies were then tested by PCR to verify the correct *mCherry* translational fusion to *rpoC*.

### Automated growth curve measurements

Cultures grown overnight of the specified microorganism were used as inocula. For fast-growing conditions, the cell suspension was diluted 1000-fold in LB or BHI and incubated at 37°C during the experiment. For slow-growing conditions, the bacterial suspension was diluted 500-fold in M9 minimal medium supplemented with 0.8% glucose, CaCl_2_ (5 mM), MgSO_4_ (32 mM), thiamine (4 μM) and micronutrient solution and incubated at 30°C. Bacterial preparations were distributed by triplicate in 96-well microplates. Growth curve experiments were performed using a TECAN Infinite 200 microplate reader (TECAN, Männedorf, Germany), with absorbance (620nm) taken at 2-minutes intervals for a period of 5 hours under continuous agitation for fast-growing conditions. For slow-growing conditions the absorbance was measured every 10 minutes for a period of 12 hours under continuous agitation. The obtained OD values were plotted against time using Microsoft Excel on logarithmic scale. The linear part of the curve was used to estimate μ from the slope with R^2^>0.995.

### Competitive fitness assays

The assays were performed as previously described^75^. Briefly, the fitness of each strain was measured relative to the Parental::*gfpmut*3* strain^15,38^ (Table S1). Each strain was cultured overnight at 37°C with 200-rpm shaking in 5 ml of LB broth or 30°C with 200-rpm shaking in 2 ml of M9 supplemented with 0.8% glucose, CaCl_2_ (5 mM), MgSO_4_ (32 mM), thiamine (4 μM) and micronutrient solution (1000X). After 1000-fold dilution in PBS, flow cytometry (FC) (BD Fortessa LSR X-20, BD Biosciences) was performed to assess the number of cells and to verify the fluorescence of *V.cholerae* (GFP)-expressing. Cultures were then mixed at a 1:1 ratio with the latter strain. Initial proportions were confirmed by FC, and mixtures were diluted 5,000-fold in fresh LB or 1,000-fold for M9 and competed for 18 h at 37°C or 24 h 30°C, respectively, with shaking at 200 rpm (∼12 and ∼10 generations, respectively). Final proportions were determined by FC as described above. The fitness of each mutant relative to the GFP-expressing *V. cholerae* strain was determined using the formula:

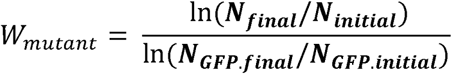

where *W_mutant_* is the fitness of the movant strain under study, *N_initial_* and *N_final_* are the quantity of the movant strain before and after the competition (nonfluorescent), respectively. *N_gfp.initial_* and *N_gfp.final_* are the numbers of cells counts of Parental::*gfpmut3*\* before and after the competition, respectively. The experiments were performed at least 3 times with 2 or more biological replicates. Relative fitness (W_rel_) is the ratio of the W of each movant strain to the W of the corresponding parental strain. The percentage of variation (ΔW%) represents the difference between the movant and the corresponding parental strain.

### Marker Frequency Analysis (MFA)

Genomic DNA was extracted from the bacterial culture when the OD_600nm_ reached ∼0.2 (early exponential phase). Libraries were sequenced by Genewiz from Azenta Life Sciences or SeqCoast Genomics on the Illumina® platforms (2□×□150 base pairs configuration, 20–30 million read depth, with ≥80% bases at Q30 quality). The resulting FastQ files were analyzed using R2R script to determine locus frequency across the genome^76^. Log_2_ frequencies every 1,000-bp window were plotted against replichore length and normalized to *ter1*. After MFA, *ori1* and *ter1* were quantified by averaging 20 kbp corresponding to *ori1* and *ter1* zones. The *rplKAJL-rpoBC* frequency was calculated by averaging panels corresponding to VC320-VC330. These values were used to calculate *rplKAJL-rpoBC, ori1* and *ori2* dosage relative to the *ter1* region. Sequence data was submitted to the GenBank Sequence Read Archive (SRA) as BioProject PRJNA124649.

### Time-lapse microscopy

To perform sequential imaging (time-lapse) the indicated strains were grown in LB to OD_600_∼0.5 and diluted 1: 300. Then, 3 μL were distributed in a LB-agar layer and sealed with VALAP resin (petroleum jelly, linoleic acid, kerosene 1:1:1). Observations were carried out with a Nikon Eclipse T2000U optical microscope equipped with a heat chamber to maintain the sample at 37°C during imaging. Automated imaging was performed every 2 minutes for 3 hours at 60X magnification. Focus was manually adjusted between acquisitions. Finally, image series were analyzed using Image J software.

### Fluorescence microscopy analysis

Cells in early exponential phase (OD_600nm_ ∼0.2) were fixed with 4% paraformaldehyde and stained with Hoechst 33342 (20µg/ml) for 15 min at room temperature. Fixed cells were washed twice with PBS and mounted on glass coverslips pretreated with poly-L-lysine. Finally, the coverslips were washed three times in PBS and once in Milli-Q water andmounted on glass slides using Fluorsave (Calbiochem).

Images were acquired using an inverted microscope Zeiss Axio Observer 7 inverted microscope equipped with a Colibri R(G/Y) B-UV LED-based illumination source. Images were taken with a Zeiss ZEN 2.3 Pro and an oil-immersion 63x Plan APO 1,4NA objective. Post-acquisition analysis was conducted using the ImageJ software package. Bacterial detection was performed with MicrobeJ^77^ (https://www.microbej.com/), which was also used to extract morphology measurements and fluorescence intensity data.

## Supporting information

Supplementary Video S1 Parental

Supplementary Video S2 movant Tnp+35

Supplementary Video S3 movant Tnp+1120

Supplementary Video S4 Tnp+479II

## Acknowledgements

We are grateful to Soledad Guidolín and Inés Marchesini for useful discussions. We thank Julieta Viglino for her technical help.

