## Supplementary Figures and Tables for "Evolutionary Constraints on RNA Polymerase Gene Positioning in the Genome of Fast-Growing Bacteria"

**Table S1.** Full list of plasmids and bacteria strains used in this study:

| Name | Relevant genotype or features | Reference |
|---|---|---|
| **Plasmids** | | |
| pMP96 | pSC101*rep*$^{TS}$ *oriT*$_{RP4}$ [*int*$_{\lambda}$-*xis*$_{\lambda}$, *int*$_{HK}$-*xis*$_{HK}$] | [1] |
| pASB9 | pCR-BluntII-Topo:: lox66-*dfrB1*-lox71 | [2] |
| pASB11 | pCR-BluntII-Topo:: lox66-Zeo-lox71 | [2] |
| pMP98 | *oriRK6 [5'lacZ-attL*$_{\lambda}$*-FRT-aph-FRT]* | This Study |
| pMP102 | *oriRK6 [loxP-cat-loxP-attR*$_{\lambda}$*-3'lacZ]* | This Study |
| pTSA29CXI | pSC101*rep*$^{TS}$ *oriT*$_{RP4}$ [*int*$_{\lambda}$-*xis*$_{\lambda}$*]* | [3] |
| pMP109 | pSC101*rep*$^{TS}$ *oriT*$_{RP4}$ *cre-flp* | Mazel Lab |
| pMP7 | pSW23T*::araC* P$_{BAD}$-*ccdB oriV*$_{R6K\gamma}$, *oriT*$_{RP4}$; [Cm$^{R}$] | [1] |
| pV073 | pMP7*::rpoBC-mcherry* for performing the *rpoBC-mcherry* translational fusion in the Parental and relocated strains | This Study |
| ***Escherichia coli*** | | |
| π3813 | *lacIq thi-1 supE44 endA1 recA1 hsdR17 gyrA462 zei-298::Tn10 ΔthyA*::(*erm-pir116*) [Em$^{R}$] | [4] |
| β3914 | β2163 *gyrA462 zei-298::Tn10* [Km$^{R}$ Em$^{R}$] | [4] |
| ***Vibrio cholerae*** | | |
| N16961*ChapRΔlacZ* | N16961::mTn*7hapR*$^{+}$ Δ*lacZ* | [1] |
| VG0052 | N16961::mTn*7hapR*$^{+}$ Δ*lacZ*, [3'*lacZ-attL*$_{\lambda}$*-FRT-aph-FRT*] (at the intergenic space VC319-VC320), [*FRT-cat-FRT-attR*$_{\lambda}$-5´ *lacZ*] (at the intergenic space VC330-VC331) flanking the region VC322 and VC329 encompassing *rplKA-JL-rpoBC locus*. | This study |
| Parental +35 | VG0052::*attB*'-lox66-*dfrB1*-lox71 inserted at the intergenic region between VC351-VC352. | This study |

| | | |
|---|---|---|
| Parental +1120 | VG0052::*attB'-lox66-dfrB1-lox71* inserted at the intergenic region VC1384-VC1385. | This study |
| Parental +35 Δ(*aph,cat*) | VG0052::*attB'-lox66-dfrB1-lox71* inserted at the intergenic region VC351-VC352. Kanamycin and chloramphenicol resistance cassettes were deleted using a flipase expressing plasmid. | This study |
| Parental +1120 Δ(*aph,cat*) | VG0052::*attB'-lox66-dfrB1-lox71* inserted at the intergenic region VC1384-1385. Kanamycin and chloramphenicol resistance cassettes were deleted using a flipase expressing plasmid. | This study |
| Parental +479II | VG0052::*attB'-lox66-dfrB1-lox71* inserted at the intergenic region between VCA0543-VCA0544. | This study |
| *rpoBC* Tnp+35 | *rplKA-JL-rpoBC* relocated next to its original location. Derived from Parental+35. | This study |
| *rpoBC* Tnp+1120 | *rplKA-JL-rpoBC* relocated near the *dif* region of chromosome 1. Derived from Parental +1120. | This study |
| *RpoBC* Tnp+479II | *rplKA-JL-rpoBC* relocated near the *dif* sequence of chromosome 2. Derived from Parental C2+479. | This study |
| Md(0;+35) | Merodiploid bearing two *rplKA-JL-rpoBC* copies. Additional copy inserted at the intergenic region between VC351-VC352. | This study |
| Md(0;+1120) | Merodiploid bearing two *rplKA-JL-rpoBC* copies. Additional copy inserted at the intergenic region between VC1384-1385. | This study |
| Md(0;+479II) | Merodiploid bearing two *rplKA-JL-rpoBC* copies. Additional copy inserted at the intergenic region between VCA0543-VCA0544. | This study |
| Md(+35;+1120) | Merodiploid bearing two *rplKA-JL-rpoBC* copies inserted at the intergenic region between VC351-VC352 and VC1384-1385. | This study |
| Md(+1120;+479II) | Merodiploid bearing two *rplKA-JL-rpoBC* copies inserted at the intergenic region between VC1384-1385 and VCA0543-VCA0544. | This study |

| | | |
|---|---|---|
| Md(*ΔrpoBC*;+35) | N16961::mTn*7hapR*$^{+}$ Δ*lacZ*, *rpIKA-JL-zeo*$^{R}$ at original position and *rpIKA-JL-rpoBC* at the intergenic region between VC351-VC352. | This study |
| Md(*ΔrpoBC*;+1120) | N16961::mTn*7hapR*$^{+}$ Δ*lacZ*, *rpIKA-JL-zeo*$^{R}$ at original position and *rpIKA-JL-rpoBC* at the intergenic region between VC1384-1385. | This study |
| Md(*ΔrpoBC*;+479II) | N16961::mTn*7hapR*$^{+}$ Δ*lacZ*, *rpIKA-JL-zeo*$^{R}$ at original position and *rpIKA-JL-rpoBC* at the intergenic region between VCA0543-VCA0544. | This study |
| Parental-mCherry | *rpoC* was translationally fused to *mcherry* by allelic exchange by delivering the pV073 suicide conjugative plasmids in the Parental +1120 Δ(*aph,cat*) strain. | This Study |
| Tnp+1120-mCherry | *rpoC* was translationally fused to *mcherry* by allelic exchange by delivering the pV073 suicide conjugative plasmids in the *rpoBC*Tnp+1120 Δ(*aph,cat*) strain. | This Study |
| Parental-GFP | gfpmut3*-Zeo$^{R}$ cassette was inserted in the intergenic region between VC0696-VC0697 in a Parental strain. | Soler-Bistué et al. 2017 |

**Table S2. The genomic localization of the rpoBC locus is conserved in *Vibrionaceae*.**

[a]representative *Vibrionaceae* species were selected among those whose genome is completely assembled. [b]The *oriC1* was coordinates were obtained from DoriC Database. Distance was taken from *oriC1* to *tuf*. [c] Distance between *rplKAJL-rpoBC* and *oriC1* was divided by the replichore length and multiplied by 100.

| ***Vibrionaceae* species[a]** | **Ref. sequence** | ***rpoBC* distance to *oriC1*(bp)[b]** | ***rpoBC* position (%replichore)[c]** |
|---|---|---|---|
| **Aliivibrio fischeri ES114** | NC_006840 | 198195 | 13.64 |
| **Aliivibrio fischeri MJ11** | NC_011184 | 183486 | 12.63 |
| **Aliivibrio salmonicida LFI1238** | NC_011312 | 212526 | 12.78 |
| ***Grimontia hollisae* strain ATCC 33564** | NZ_CP014056 | 179533 | 11.17 |
| ***Photobacterium profundum* SS9** | NC_006370 | 192378 | 9.42 |
| ***Vibrio alginolyticus*** | NC_022349.1 | 185156 | 11.03 |
| ***Vibrio alginolyticus* NBRC 15630 = ATCC 17749** | NC_022349 | 185156 | 11.11 |
| ***Vibrio anguillarum* 772** | NC_015633 | 233500 | 15.24 |
| ***Vibrio antiquarius*** | NC_013456 | 161726 | 9.92 |
| ***Vibrio breoganii* strain FF50** | NZ_CP016177 | 119548 | 8.52 |
| ***Vibrio campbellii* ATCC BAA-1116** | NC_022269 | 178347 | 9.52 |
| ***Vibrio cholerae* M66-2** | NC_012578 | 320154 | 22.14 |
| ***Vibrio cholerae* MJ-1236** | NC_012668 | 213700 | 13.49 |
| ***Vibrio cholerae* N16961** | NC_002505 | 335428 | 22.66 |
| ***Vibrio cholerae* O395** | NC_009457 | 208455 | 13.79 |
| ***Vibrio coralliitycus*** | NZ_CP009264 | 233616 | 13.49 |
| ***Vibrio furnissii* NCTC 11218** | NC_016602 | 294200 | 17.78 |
| ***Vibrio gazogenes* strain ATCC** | NZ_CP018835 | 351730 | 20.27 |
| ***Vibrio harveyi* ATCC BAA-1116** | NC_009783 | 181696 | 9.65 |
| ***Vibrio mimicus*** | NZ_CP016383 | 226812 | 14.12 |
| ***Vibrio natriegens* NBRC 15636 = ATCC 14048 = DSM 759** | NZ_CP009977 | 339317 | 20.89 |
| ***Vibrio nigripulchritudo* SFn1** | NC_022528 | 753742 | 36.68 |
| ***Vibrio parahaemolyticus* RIMD 2210633** | NC_004603 | 167563 | 10.11 |

| | | | |
|---|---|---|---|
| ***Vibrio parahaemolyticus* RIMD 2210633** | NC_004603.1 | 167449 | 10.18 |
| ***Vibrio scophthalmi* strain VS-12** | NZ_CP016307 | 295263 | 18.08 |
| ***Vibrio* sp. Strain EJY3** | NC_016613 | 158706 | 9.13 |
| ***Vibrio splendidus* LGP32** | NC_011753 | 173660 | 10.53 |
| ***Vibrio tubiashii* ATCC 19109** | NZ_CP009354 | 289887 | 17.6 |
| ***Vibrio* vulnificus CMCP6** | NC_004459 | 170134 | 10.37 |
| ***Vibrio vulnificus* MO6-24/O** | NC_014965 | 169666 | 10.62 |
| ***Vibrio vulnificus* YJ016** | NC_005139 | 111959 | 6.68 |

**Table S3.** Generation time (GT) of *rplKA-JL-rpoBC movants* generated in this study determined by automated growth curves. Results are shown as mean ± sd, with n>6.

| STRAIN | GROWTH CONDITION | | |
|---|---|---|---|
| | LB | BHI | M9 1% Glucose |
| **Parental +35** | 16.784 ± 0.230 | 16.563 ± 0.139 | 80.096 ± 3.336 |
| **Parental +1120** | 16.758 ± 0.450 | 16.550 ± 0.384 | 81.518 ± 4.790 |
| **Parental +1120 Δ(*aph,cat*)** | - | 17.009 ± 0.462 | - |
| **Parental +479 II** | 17.478 ± 0.839 | 16.977 ± 0.695 | 74.813 ± 7.410 |
| ***rpoBC* tnp+35** | 16.717± 0.658 | 16.709 ± 0.463 | 73.382 ± 3.716 |
| ***rpoBC* tnp+1120** | 18.263 ± 0.460 | 19.139 ± 0.726 | 73.823 ± 7.286 |
| ***rpoBC* tnp+479 II** | 19.480 ± 0.895 | 18.908 ± 1.190 | 70.483 ± 13.563 |
| **Md(0;+35)** | - | 17.262 ± 0.883 | 70.631 ± 3.480 |
| **Md(0;+1120)** | - | 17.404 ± 0.625 | - |
| **Md(0;+479II)** | - | 18.402 ± 0.620 | - |
| **Md(+35;+1120)** | - | 17.882 ± 0.547 | 79.742 ± 8.505 |
| **Md(+1120;+479II)** | - | 18.618 ± 0.784 | 86.056 ± 12.877 |
| **Md(Δ*rpobc*;+35)** | - | 17.814 ± 0.516 | - |
| **Md(Δ*rpobc*;+1120)** | - | 18.740 ± 0.774 | - |
| **Md(Δ*rpobc*;+479II)** | - | 18.614 ± 0.686 | - |
| **Tnp+1120 Δ(aph,cat)** | - | 18.146 ± 0.469 | - |
| **Parental *rpoC-mCherry*** | - | 16.958 ± 0.527 | - |
| **Tnp+1120 *rpoC-mCherry*** | - | 18.499 ± 0.798 | - |

**Table S4.** Absolute fitness (W) of *rplKA-JL-rpoBC* movants generated in this study measured by pairwise competition against the parental strain carrying *gfpmut3**. Results are shown as mean ± sd, with n>6.

| **STRAIN** | **GROWTH CONDITION** | | |
|---|---|---|---|
| | LB | BHI | M9 1% Glucose |
| **Parental +35** | 1.188 ± 0.0671 | 1.183 ± 0.054 | 1.321 ± 0.182 |
| **Parental +1120** | 1.206 ± 0.066 | 1.186 ± 0.083 | 1.175 ± 0.041 |
| **Parental +479 II** | 1.179 ± 0.081 | 1.142 ± 0.055 | 1.292 ± 0.107 |
| ***rpoBC* tnp+35** | 1.180 ± 0.055 | 1.200 ± 0.062 | 1.290 ± 0.146 |
| ***rpoBC* tnp+1120** | 1.010 ± 0.038 | 1.073 ± 0.040 | 1.138 ± 0.053 |
| ***rpoBC* tnp+479 II** | 1.101 ± 0.076 | 1.076 ± 0.052 | 1.243 ± 0.118 |
| **Md(0;+35)** | - | 1.169 ± 0.041 | - |
| **Md(Δ*rpoBC*;+35)** | - | 1.127 ± 0.030 | - |
| **Md(0;+1120)** | - | 1.185 ± 0.085 | - |
| **Md(Δ*rpoBC*;+1120)** | - | 1.121 ± 0.062 | - |

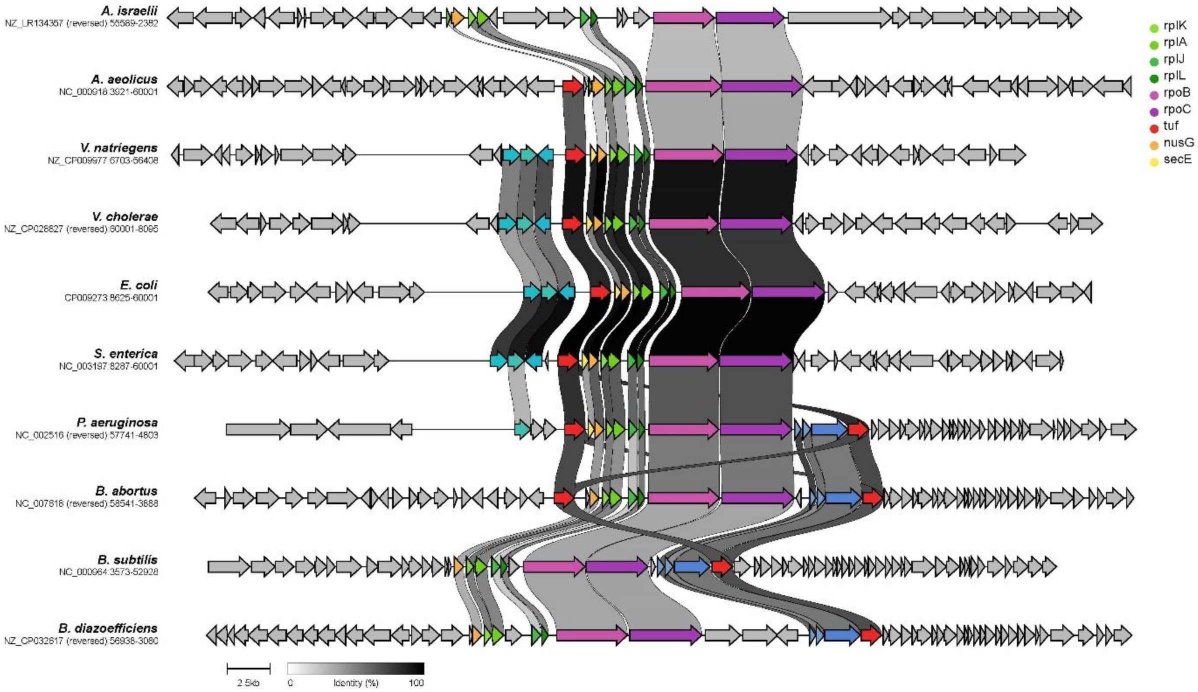


**Figure S1. The *rplKAJL-rpoBC* locus is conserved in bacteria.** Gene clustering of the *rplKAJL-rpoBC* locus was performed using the Clinker[5] free tool directly from Genebank sequence files. Links are drawn between similar genes in neighbouring clusters and shaded based on sequence identity (0% white, 100% black).

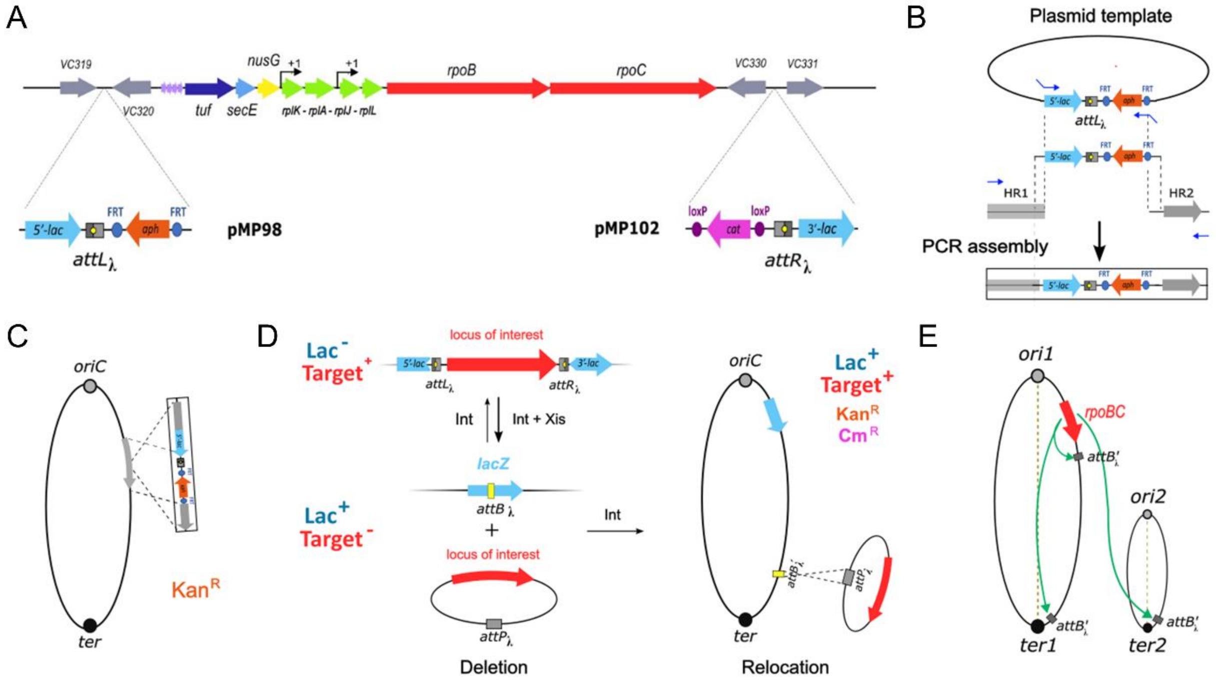


**Figure S2: *rplKAJL-rpoBC* relocation using lamboid phage recombination. (A)** Detailed view of *rplKAJL-rpoBC* locus. Thick arrows indicate genes colored according function (Green, translation; light blue , secretion; tRNAs in purple, dark blue translation-transcription coordination; red transcription; and grey, other). The dotted lines indicates the insertion of *attL* and *attR* sites respectively and the template plasmid coming from. **(B)** DNA assembly of selected *att* (grey) site linked to a kanamycin resistance gene (orange) and flanked by two homology regions (HR). (**C)** The *att*-fragment is targeted to its genomic location by double cross and antibiotic selection. (**D)** Upon transtient expression of integrase (Int) and excisase (Xis), the excisive recombination reaction leads to deletion of *rplKAJL-rpoBC* (red). **(E)**The insertion of an *attB* (attB') elsewhere permits *attP×attB'* crossover that reinserts the locus of interest. The *attB'* orientation determines in which DNA strands the insertion will occur.

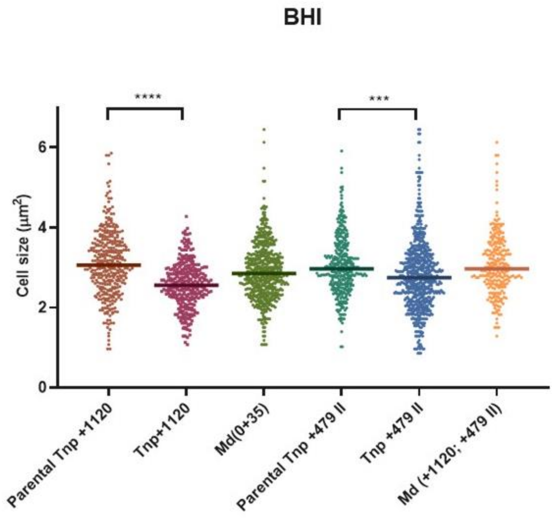


**Figure S3: *rpoBC* relocation causes a cell length in fast growing conditions.** The indicated strains were cultured in BHI until early exponential phase ($OD_{450mn}$≈0.2). Then microscopy images were taken and cells were measured using Image J. The length of at least 100 cells of each strain was measured. Overall, the length of Tnp+1120 and TnpII+479 was significantly lower than their parental strain when cultured in BHI.

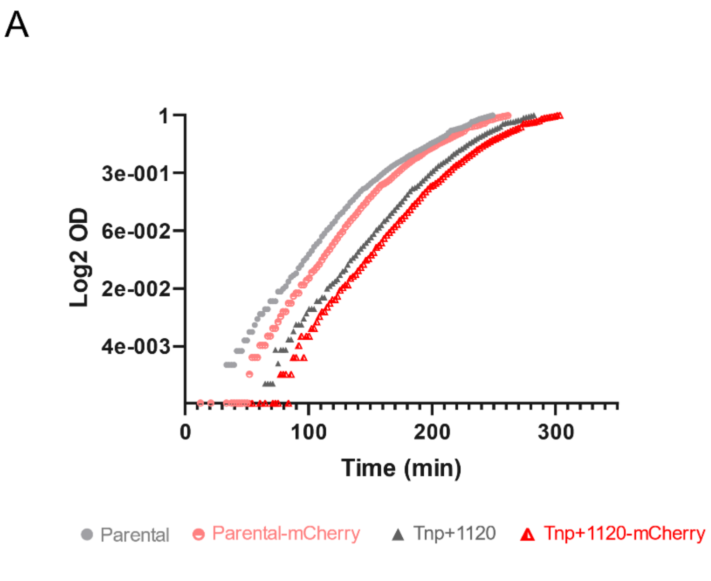


**Figure S4. RNAP β´ subunit tag did not affect the phenotypes related to *rplKA-JL-rpoBC* genomic position.** (A) The growth rate of parental and movant strains were determined in automated growth curves performed in BHI media at 37°C. The left panel shows a representative growth curve. (B) The effect of the translational fusion of RpoC and mCherry on TG was quantified by averaging the difference in TG of each motile relative to its parent in at least 3 independent experiments. Results are expressed as percentage variation (%ΔGT) with 95% CI with respect to the parental strains. The percentage variation with.Statistical significance was analysed by two-tailed t-test (ns. not significant;p>0.05; *. p<0.05; **. p<0.01; ***. p<0.001;****. p<0.0001).

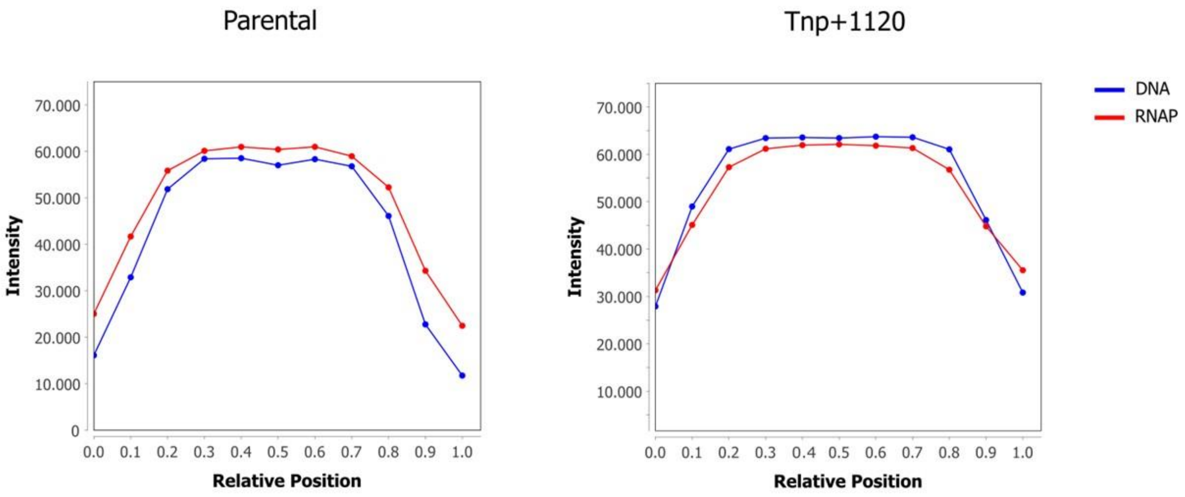


**Figure S5. The mCherry signal closely followed the DNA signal in Parental and movant strain.** Profiles of relative fluorescent intensities of DNA (blue) and RpoC (red) in the Parental-mCherry (left) and movant Tnp+1120-mCherry (right) analized with MicrobeJ[6] plugin (n=400).

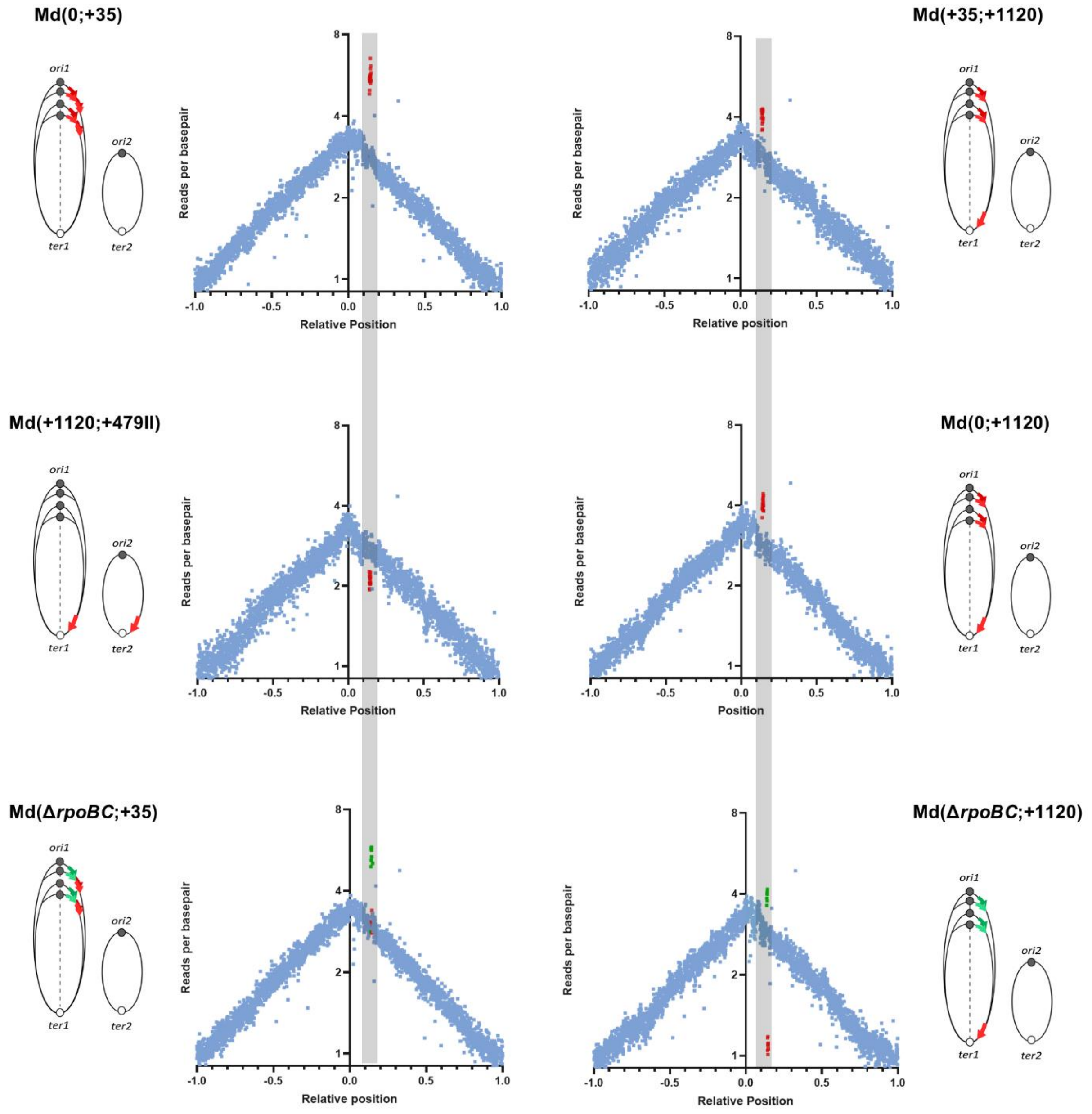


**Figure S6. Gene dosage of genes forming the *rplKAJL-rpoBC* locus in merodiploid and Δ*rpoBC* strains.** Marker Frequency Analysis (MFA) of exponentially growing strains using a corrected reference sequence of Chr1. The log2 of the number of reads starting at each base is plotted against their relative position on Chr1. The position of *ori1* is set to 0 to better represent bidirectional replication. Dots indicate the average of 1000 bp windows. Red dots correspond to *rplKAJL-rpoBC* genes in Md (0;+35). Md(0;+1120). Md(+35;1120) and Md(+1120;+479II). In Md(Δ*rpoBC;+35)* and Md(Δ*rpoBC*;+1120) green dots represent *rplKAJL* genes and red dots indicates *rpoBC* genes. Ovals represent the two *V. cholerae* chromosomes and red arrows indicate the *rplKA-JL-rpoBC* locus position and green arrows represent *rplKAJL-zeo* position.

**Table S5.** Sequences of phage lambda site-specific recombination sites

| ***att* Site** | **Sequence** |
|---|---|
| *attL* | TTTATACTAAGTTGGCATTATAAAAAAGCATTGCTTATCAATTTGTTGCAACGAACAGGTCACTATCAGTCAAAATAAAATCATTATT |
| *attR* | GTATAAAAAAGCTGAACGAGAAACGTAAAATGATATAAATATCAATATATTAAATTAGATTTTGCATAAAAAACAGACTACATAATACTGTAAAACACAACATATGCAGTCACTATGAATCAACTACTTAGATGGTATTAGTGACCT |
| *attB'* | GGAAGCCTGCTTTTTTATACTAACTTGAGCGAC |

**Supporting Material References:**
